# Quantitative occupancy of myriad transcription factors from one DNase experiment enables efficient comparisons across conditions

**DOI:** 10.1101/2020.06.28.171587

**Authors:** Kaixuan Luo, Jianling Zhong, Alexias Safi, Linda K. Hong, Alok K. Tewari, Lingyun Song, Timothy E. Reddy, Li Ma, Gregory E. Crawford, Alexander J. Hartemink

## Abstract

Over a thousand different transcription factors (TFs) bind with varying occupancy across the human genome. Chromatin immunoprecipitation (ChIP) can assay occupancy genome-wide, but only one TF at a time, limiting our ability to comprehensively observe the TF occupancy landscape, let alone quantify how it changes across conditions. We developed TOP, a Bayesian hierarchical regression framework, to profile genome-wide quantitative occupancy of numerous TFs using data from a single DNase-seq experiment. TOP is supervised, and its hierarchical structure allows it to predict the occupancy of any sequence-specific TF, even those never assayed with ChIP. We used TOP to profile the quantitative occupancy of nearly 1500 human TF motifs, and examined how their occupancies changed genome-wide in multiple contexts: across 178 cell types, over 12 hours of exposure to different hormones, and across the genetic backgrounds of 70 individuals. TOP enables cost-effective exploration of quantitative changes in the landscape of TF binding.

## Introduction

Genes are expressed differently in different types of cells and under different conditions. This response of a cell’s gene expression to its internal and external context is enacted in large part through the tuned occupancy of transcription factors (TFs) across the genome. To understand how TFs regulate gene expression, it is critical to determine how likely they are to be present at each location in the genome over time, and how that likelihood changes across varying genetic backgrounds, different cell types, and dynamic environmental conditions. We can measure the quantitative occupancy of one TF along the genome using chromatin immunoprecipitation followed by high-throughput sequencing (ChIP-seq), provided that a selective antibody exists for the TF. While the ENCODE consortium has generated such data for more than 100 human TFs, the data are typically from only a small number of cell types because of a major limitation of ChIP-seq: a separate experiment is required for each TF in each cell type under each condition. Profiling the time-varying genome-wide occupancy of a large set of TFs across a broad range of cell types and conditions is currently impractical since it would require thousands of antibodies and millions of separate ChIP experiments.

An alternative strategy for profiling genome-wide TF occupancy is to exploit DNase-seq or ATAC-seq data, which many groups and consortia have generated for a large number of cell types and experimental conditions^1–4^. The primary advantage of this strategy is that a single DNase-seq or ATAC-seq experiment can be used to profile the occupancy of many different TFs at once, and a number of methods employing this strategy have been proposed in recent years^5–17^.

Although multiple methods have been developed to predict TF binding (see Supp. Table S1 for an overview of the modeling frameworks used by a number of these methods), many of them require data types beyond DNase^14–17^, making them less efficient at profiling TF occupancy across multiple cell types, conditions, or individuals with different genetic backgrounds than methods requiring only one data type. Furthermore, most existing methods model TF occupancy in a binary fashion—each TF is simply considered present or absent at each location in the genome—ignoring the wealth of quantitative information available in the data^18^. While this modeling assumption is consistent with the pervasive practice of binary peak-calling in high-throughput sequencing data, it is inconsistent with our knowledge that at different genomic locations, TFs exhibit different levels of occupancy (likelihood of being bound at that location across the cells in a population) in accordance with prevailing energetic and thermodynamic conditions, including competition with other TFs and nucleosomes^2,19–23^. It is also inconsistent with growing evidence that quantitative levels of TF occupancy can play a significant role in regulating gene expression^24–27^. Therefore, it is important that statistical models be developed with a quantitative perspective, allowing us to monitor subtle changes in TF occupancy over time across different genetic backgrounds, cell types, and conditions.

Here, we describe a novel method called TF Occupancy Profiler (TOP) that integrates DNase-seq data with information about TF binding specificity (in our case, PWM motifs) to predict the quantitative occupancy of multiple TFs genome-wide. In contrast to earlier methods like CENTIPEDE ^5^, PIQ^10^, and msCentipede^11^, TOP is supervised, meaning we can use available ChIP-seq data to train it to high accuracy. Importantly, and in contrast to earlier methods like MILLIPEDE ^6^ and BinDNase^12^, TOP employs a Bayesian hierarchical regression framework, which allows it to obtain both TF-specific and TF-generic model parameters by borrowing information across the full spectrum of training TFs and cell types. The hierarchical nature of TOP is significant because it enables us to predict the occupancy of TFs for which we lack training data, including ones that have never before been profiled with ChIP.

We used TOP to predict the genome-wide quantitative occupancy of ∼1500 TF motifs across 178 human cell types, increasing our cell-type–specific view of TF occupancy in human cells over 200-fold relative to the ENCODE ChIP-seq data used to train the model. We have made these predicted TF occupancy profiles freely available for the community. We used them to construct a cell-type specificity map for different TFs, and identified TFs with selective binding and differential occupancy across cell types. To demonstrate TOP’s ability to elucidate the dynamics of TF occupancy, we collected DNase-seq data from A549 cells at 12 time points over 12 hours of glucocorticoid exposure^28^ and used TOP to efficiently screen nearly 1500 TF motifs for increased or decreased occupancy throughout the genome following treatment; we did follow-up ChIP experiments for six of those factors to validate our predictions. We show similar results in separate cells stimulated with androgen or estrogen, two other steroid hormones that act through closely related mechanisms. In another application, we predicted quantitative TF occupancy for the same ∼1500 TF motifs across 70 Yoruba lymphoblastoid cell lines (LCLs), and mapped thousands of genetic variants associated with quantitative TF occupancy across individuals (which we term ‘topQTLs’). These topQTLs suggest specific mechanistic explanations for the functional impact of genetic variants within regulatory regions. In summary, TOP offers a cost-effective strategy for profiling the occupancy of multiple TFs in a single experiment, markedly enhancing our ability to explore subtle quantitative changes in TF occupancy across cell types, conditions, and genetic variants.

## Results

### Bayesian hierarchical regression accurately predicts quantitative TF occupancy from DNase-seq data

Training TOP entailed two basic steps, as illustrated in Fig. 1. First, we used motif matches to enumerate candidate binding sites, and extracted DNase and ChIP data centered on each site for training. Second, we used MCMC to fit our Bayesian hierarchical regression model on spatially-binned DNase data. Owing to its hierarchical nature, once TOP is trained, we can use it to predict occupancy for any TF in any cell type or condition for which we have DNase-seq data, regardless of whether any ChIP-seq data have ever been collected for that TF.

**Fig. 1.**
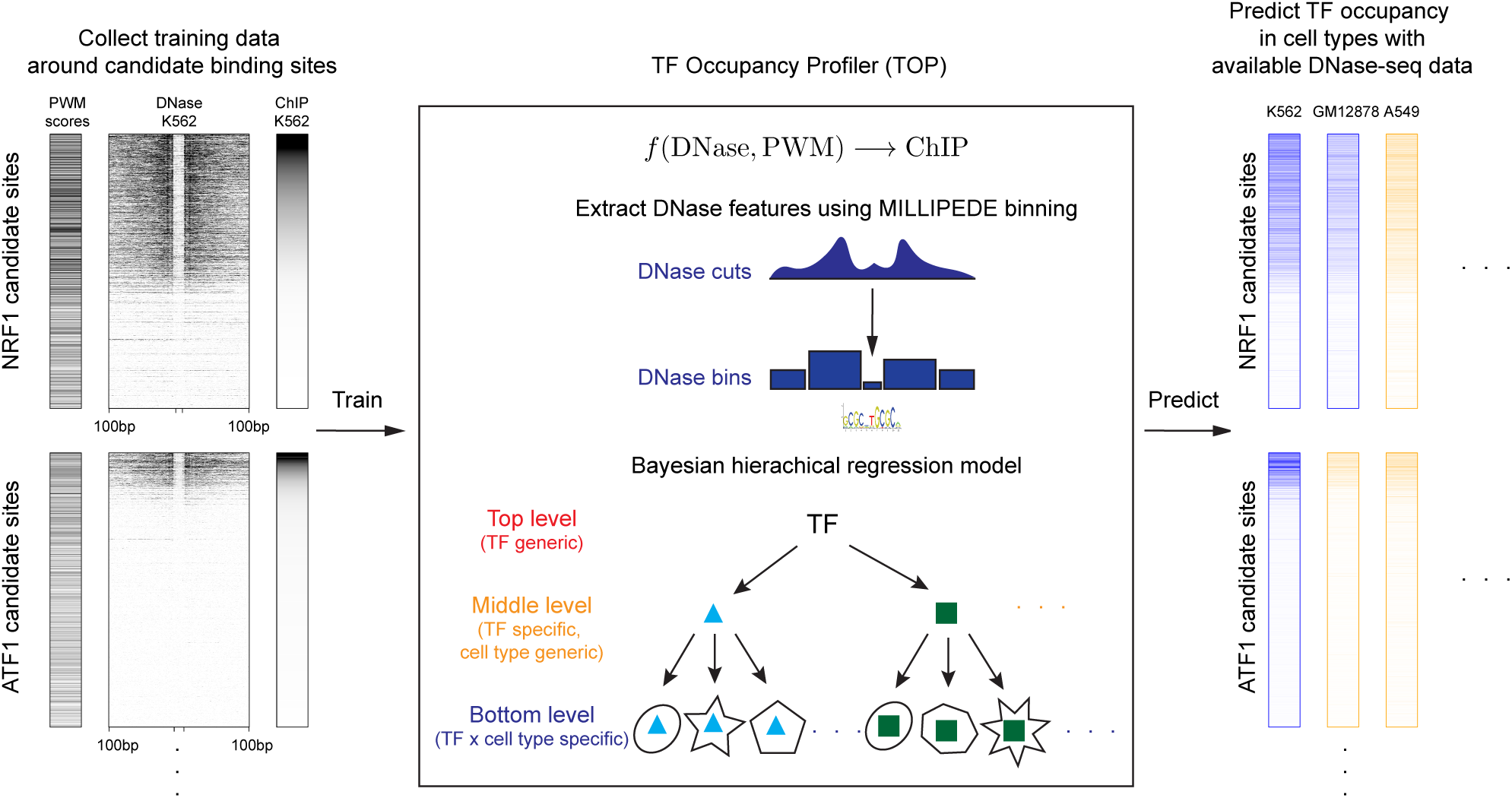
Schematic outline of the TF Occupancy Profiler (TOP) workflow. (Left) Collect training data. For a sequence-specific TF with a known PWM, compute its candidate binding sites throughout the genome. Then, around each of those sites, collect ChIP-seq and DNase-seq data from the same cell type. (Center) Extract DNase features using MILLIPEDE binning and fit a Bayesian hierarchical regression model to the training data. Bottom level models in the hierarchy make predictions in a TF × cell-type– specific manner, middle level models extend prediction in a TF-specific manner to new cell types, and the top level model extends prediction in a TF-generic manner to new TFs. (Right) Predict occupancy for TFs across cell types. Blue columns indicate a cell type where ChIP-seq measurements are available, allowing us to evaluate the predictive accuracy of our bottom level models. Orange columns indicate a cell type in which we make novel predictions of TF occupancy using middle level parameters of the hierarchical model.

We predicted the quantitative number of ChIP-seq reads around candidate TF binding sites using a site-centric approach, as employed by CENTIPEDE ^5^ and its successors. Specifically, for each TF, we first identified candidate binding sites by motif scanning with a permissive threshold (using FIMO with *P*-value < 10^−5^)^29^. Then, for each cell type, we considered Dnase cleavage events occurring within 100 bp of the candidate binding site. Similarly, we counted the number of ChIP-seq reads within 100 bp of the candidate binding site to serve as the target of our regression when training TOP. Both DNase and ChIP-seq counts were normalized by library size to account for differences in sequencing depth. We simplified the DNase data into five predictive features using bins that aggregate the number of cleavage events occurring within the motif itself, as well as within two non-overlapping flanking regions upstream and downstream; this is the same binning scheme used in the MILLIPEDE model^6^, and markedly reduces the potential impact of DNase digestion bias^6,8,11,30^.

As an alternative, we tried extracting DNase features using wavelet-transformed multi-scale signals from coarse to fine spatial resolution. However, after variable selection using LASSO, we found only the coarsest resolutions yielded significant features for predicting TF occupancy, while fine resolution features were essentially irrelevant (Supp. Fig. S1). Moreover, the simpler MILLIPEDE binning scheme achieved comparable or better prediction accuracy than optimally selected wavelet features (Supp. Fig. S2). As an added benefit, when fitting TOP to a large number of different TFs across many diverse cell types, the five-bin scheme demonstrated superior computational efficiency and better generality in capturing common features across TFs and cell types. Thus, the results that follow are all based on DNase data aggregated into five bins.

We chose to use a Bayesian hierarchical model because it allows statistical information to be borrowed across TFs and cell types. TOP’s hierarchical structure has three levels (Supp. Fig. S3). The bottom level of the hierarchy contains model parameters specific for each TF × cell-type combination for which ChIP-seq training data is available. In the middle level, one set of TF-specific but cell-type–generic model parameters is shared across all training cell types for each TF. Finally, the top level has one set of TF-generic parameters jointly learned from all TFs. In other words, we obtain more general model parameters as we move to higher levels of the hierarchy. Once TOP’s parameters have been trained, we can predict occupancy for any TF in any cell type in which we have collected DNase-seq data by using a model from the appropriate level of the hierarchy.

We evaluated TOP’s performance in terms of its fit to quantitative TF occupancy as measured experimentally by ChIP (Fig. 2). TOP predicted quantitative occupancy with varying degrees of accuracy across different TFs (Figs. 2a and 3). In light of technical differences and possible batch effects between DNase-seq data generated in different ENCODE labs, we trained two separate hierarchical models for data from Duke and from Washington (UW), achieving comparable performance between them (Figs. 2b and 3). In general, while bottom level models achieved the highest prediction accuracy (median correlation of 0.70 for Duke and 0.75 for UW), middle level models performed equally well (0.70 and 0.75), and top level models performed nearly as well (0.68 and 0.74) (Fig. 2b). This indicates that for a TF that has been profiled with ChIP in some cell type, we can use the TF’s middle level model to predict its occupancy in any other cell type with available DNase data. In addition, for TFs that have never been profiled with ChIP, the top level TF-generic model will still tend to provide good predictions of quantitative occupancy. Our predicted occupancy accurately matched quantitative ChIP-seq occupancy in various cell types, and allowed us to explore TF occupancy in cell types like the embryonic stem cell line H9ES in which no TF ChIP data have been reported (Fig. 2c). The quantitative predictions produce composite landscapes that sensitively reflect cell-type–specific changes in TF occupancy.

**Fig. 2.**
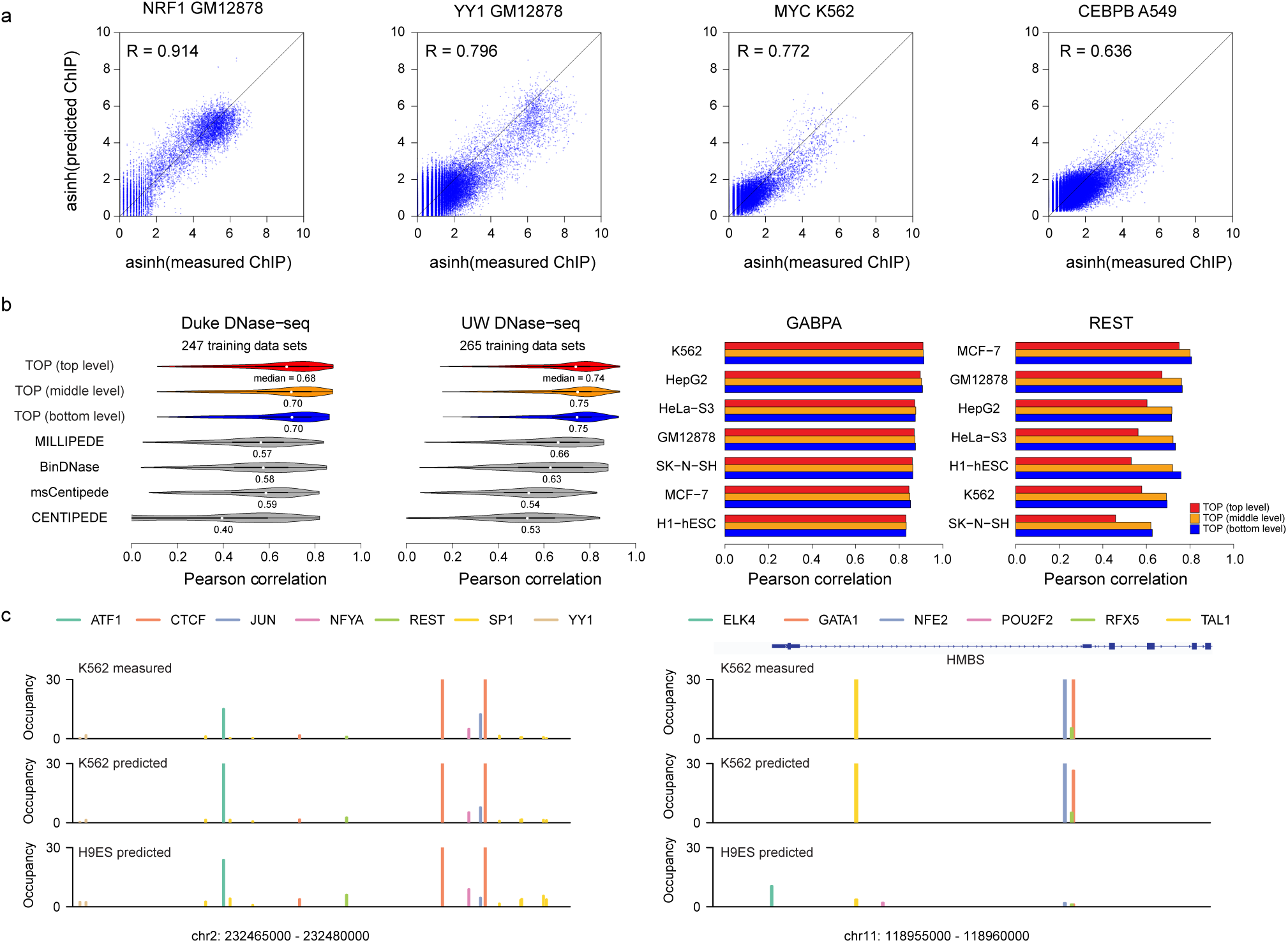
Evaluation of TOP results. (**a**) Scatter plots show predicted vs. measured occupancy of a specific TF in a specific cell type, with dots representing the candidate binding sites across the genome. The four examples are chosen to represent a range of model performance, from better to worse. (**b**) Left: Separately for Duke and UW DNase data, violin plots show distribution of Pearson correlations between predicted and measured TF occupancy (sqrt transformed) across TFs and cell types. Predictions were made with TOP models at each level of the hierarchy, in comparison with CENTIPEDE, msCentipede, MILLIPEDE, and BinDNase (using log odds of binding probability as a quantitative measurement of TF occupancy). Right: For many TFs, GABPA being one, all three levels exhibited similar correlation across various cell types. In contrast, for a few TFs, REST being one, the top level model performed markedly worse than bottom and middle level models, suggesting that TF-specific parameters enable more accurate prediction in such cases. (**c**) Predicted TF occupancy landscapes for two genomic regions in K562 and H9ES cell types. For K562, ChIP-seq data for these TFs are available and are displayed for comparison; for H9ES, no ChIP-seq data are available so TOP provides a novel view of TF occupancy in this embryonic stem cell line. Left: an example genomic region where the occupancy landscape did not change markedly between K562 and H9ES. Right: an example genomic region near the HMBS gene (involved in heme biosynthesis) where GATA1, TAL1, and NFE2 exhibited clear cell-type–specific occupancy.

To compare with alternative existing methods, since our goal is to efficiently and accurately predict quantitative TF occupancy for candidate binding sites using only a single DNase experiment, we focused our comparisons on CENTIPEDE^5^, msCentipede^11^, MILLIPEDE^6^, and BinDNase^12^ (see Methods for a discussion of why these were chosen). These predict TF binding in a site-centric framework but generate only predicted TF binding probabilities rather than ChIP-seq read counts. However, since the CENTIPEDE paper showed a substantial correlation between its TF binding predictions (posterior log odds) and ChIP-seq read counts (sqrt transformed), we used the posterior log odds of TF binding as a proxy for quantitative ChIP-seq predictions. Our results indicate TOP achieves significantly greater accuracy than all four of the other methods on both Duke and UW DNase data (Fig. 2b).

### TOP reveals a spectrum of predictability across TFs and cell types

Across TFs, we observed a spectrum of predictability of TF occupancy, as indicated by the blue squares in Fig. 3. Predictability was correlated with the degree of DNase depletion at the motif (Supp. Fig. S4). For TFs with higher prediction accuracy, like NRF1 and ATF1, we observed clear profiles of depletion within motif regions and elevation at nearby flanking regions (Supp. Fig. S5), suggesting direct TF–DNA contact. Many of these TFs have previously been classified as pioneer factors^10^. In contrast, TFs with lower prediction accuracy in the ENCODE data, like STATs and SREBPs, showed less marked elevation at nearby flanking regions, and weak or no depletion at motif regions (Supp. Fig. S5). Weaker DNase depletion profiles may result from transient binding with short residence time—known to occur with nuclear receptors and the AP-1 complex^23,30–32^—or from ChIP data that include many indirect binding events. For some TFs, we observed a high prediction accuracy in most cell types, but a lower prediction accuracy in just one or two cell types. DNase profiles in the latter cases exhibited markedly weaker depletion (Supp. Fig. S6).

**Fig. 3.**
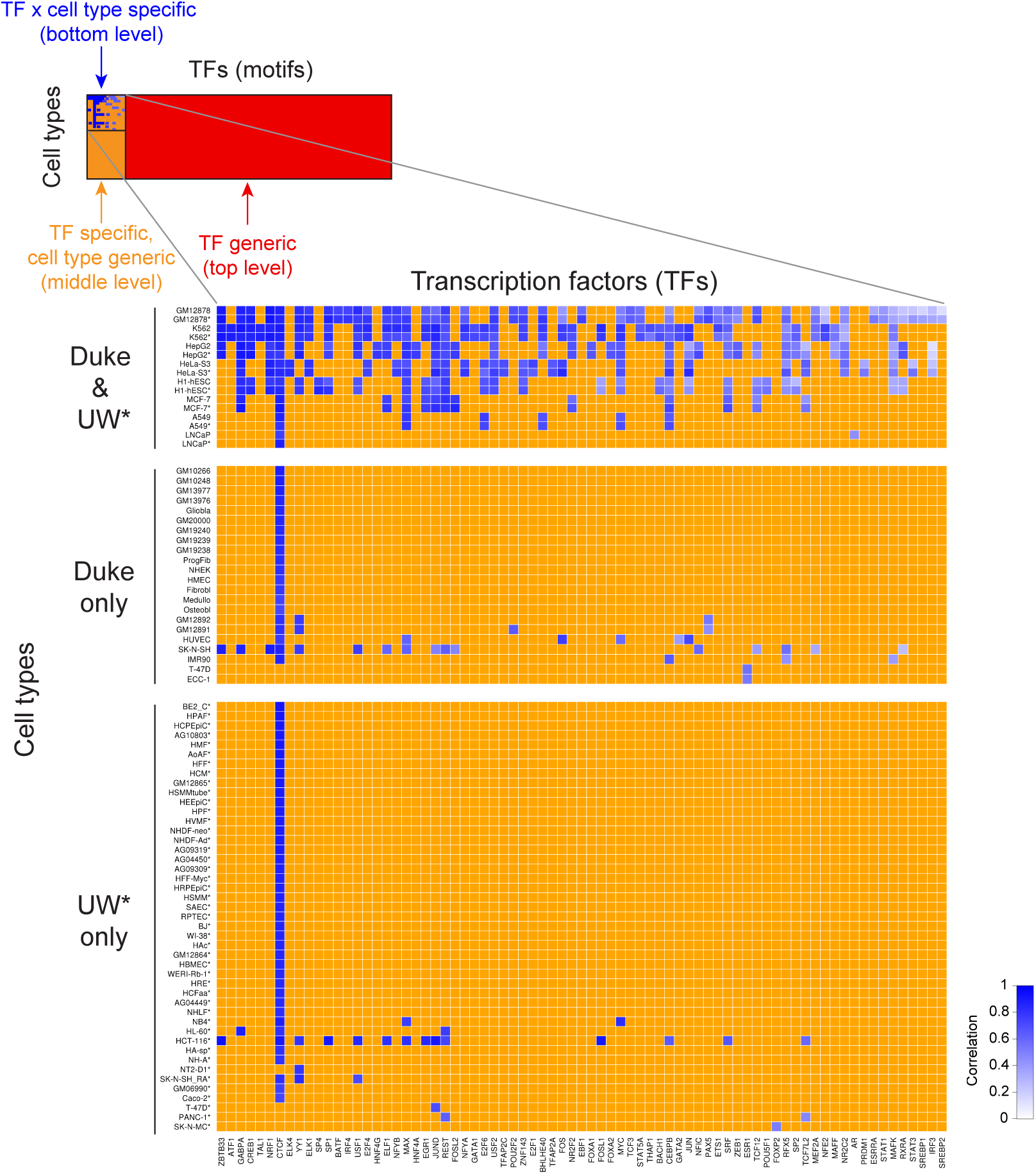
TOP provides quantitative TF occupancy profiles for nearly 1500 TF motifs in 178 cell types. (Top left) Blue squares represent the TF × cell-type combinations profiled with ChIP-seq as part of the ENCODE project. For each of these TFs, we used a middle level (TF-specific, cell-type–generic) TOP model to generate new occupancy predictions across the rest of the 178 cell types (orange squares). We then used a top level (TF-generic) model to generate new occupancy predictions for the remainder of the 1496 TFs (red squares). (Zoomed inset) In the case of TF × cell-type combinations with ChIP-seq data, we computed the accuracy of TOP predictions; shades of blue indicate the correlation between predicted and measured occupancy. In this submatrix, columns (TFs) and rows within each block (cell types) were sorted by average accuracy, revealing a spectrum of predictability. TFs toward the left were on average more predictable, while TFs to the right were less. Row order is less informative because, except in the top block, it was mainly driven by trivial fluctuations in the predictability of CTCF (in most cell types, CTCF is the only factor whose occupancy was profiled by ENCODE).

TOP uses PWM scores to provide information about which sites are more or less likely to be bound in any cell type or condition. However, in the absence of genetic variation, the PWM score of a particular site does not change across cell types or conditions, so TOP’s ability to quantify changes in TF occupancy in such situations depends entirely on changes in the DNase data. As expected, when we compared them as single features, the overall level of DNase cleavage was almost always more correlated with ChIP-seq occupancy across cell types than was the PWM score (Supp. Fig. S7).

Having established the reliability of TOP’s predictions, we applied it to data from different contexts to illustrate the biological insights that arise from its ability to efficiently compare quantitative occupancy for myriad TFs across conditions; each of the remaining three subsections explores one of these applications: changes in TF occupancy across different cell types, in response to dynamic environmental conditions, and in the context of genetic variation.

### TOP maps out the cell-type specificity of TF occupancy

TFs regulate gene expression in a cell-type–specific manner. To assess TF occupancy differences across cell types, we constructed a cell-type differential occupancy map for multiple TFs to reveal distinct patterns in how TFs direct the gene regulation programs of different cell types. For each TF, we calculated the percentage of candidate sites in each cell type showing occupancy significantly higher or lower than the mean across cell types (FDR < 10%); we then clustered TFs on the basis of this measure of cell-type specificity (Fig. 4a). Some TFs—including TAL1, GATA1, and NRF1—displayed large differences in occupancy among cell types, whereas the occupancy of other TFs—like the SPs—was quite cell-type–invariant (Fig. 4b). Lending credence to these results, we successfully recovered TFs known to be specifically or differentially expressed in certain cell types. For instance, as expected, we saw that POU5F1 (also known as Oct4) occupancy was significantly higher in stem cells, HNFs (hepatocyte nuclear factors) were higher in liver cells, GATAs were higher in K562, REST was lower in medulloblastoma, etc.

**Fig. 4.**
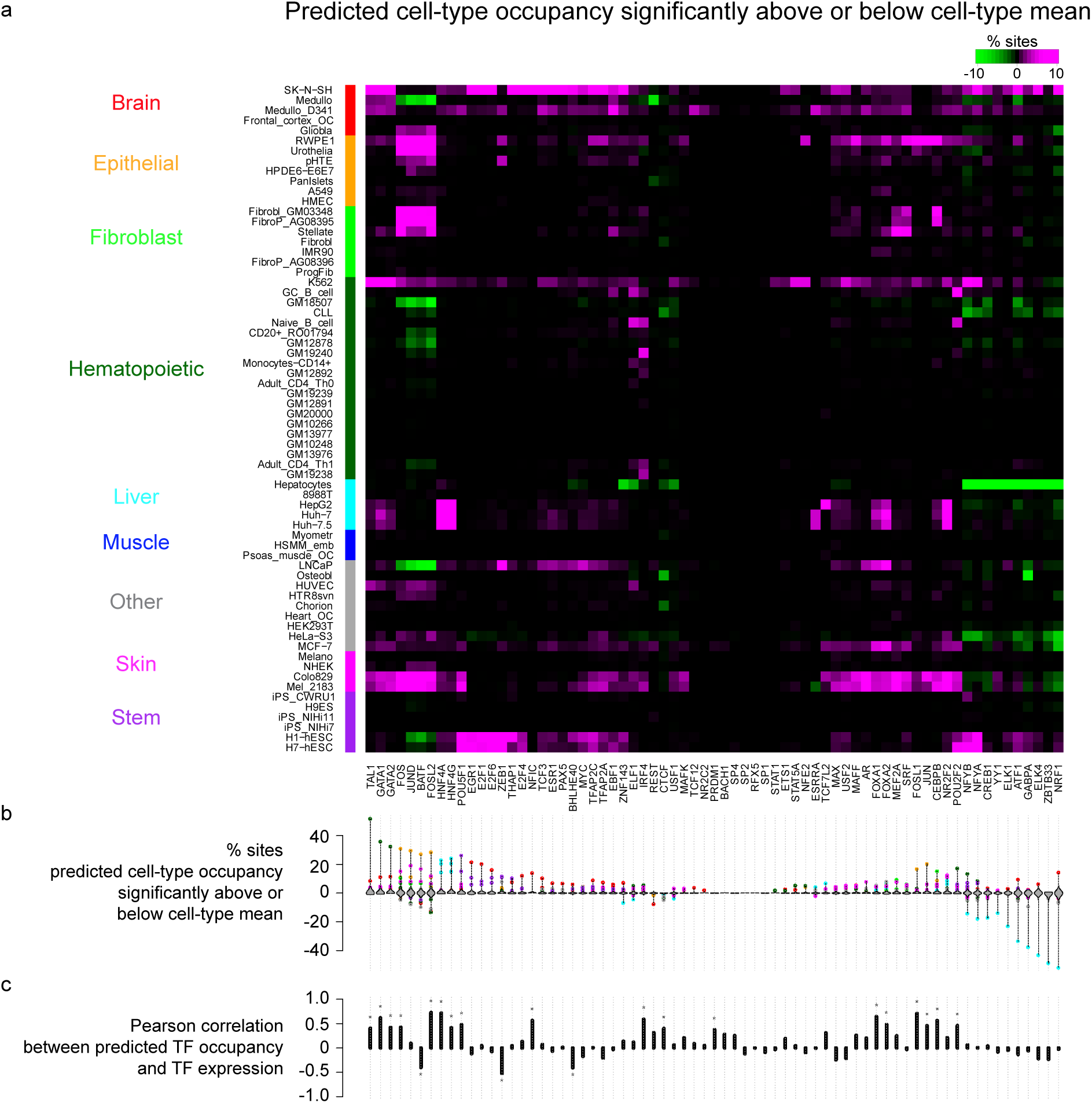
Cell-type specificity matrix of TFs. (**a**) Percentage of sites with predicted cell-type occupancy significantly above or below predicted cell-type mean occupancy (FDR < 10%). Cell types were first grouped by lineage (ordered alphabetically) ^58^, and within each lineage group were ordered by hierarchical clustering. TFs were ordered by hierarchical clustering (with optimal leaf ordering ^59^). (**b**) Violin plots show for each TF the distribution across cell types of the percentage of sites exhibiting significantly differential occupancy. Colored dots highlight cell types with at least 3% of sites exhibiting significantly differential occupancy (color reflects lineage of cell type; for instance, liver and brain exhibit frequent differential occupancy). (**c**) Pearson correlation across cell types between average predicted occupancy and gene expression of each TF (in this plot, we used only Duke DNase data because corresponding gene expression was measured in each of the cell types).

To explore the relationship between a TF’s concentration (here approximated by its gene expression level) and its occupancy, we computed the correlation between each TF’s average level of occupancy in each cell type with its gene expression level in that same cell type, and observed several categories of TFs with different relationships (Fig. 4c). Many TFs showed significant positive correlations between their gene expression level and average occupancy, most of which are known to be cell-type–specific TFs, such as FOSL2, HNF4A, FOXA1, and POU5F1 (Supp. Fig. S8a). Surprisingly, three TFs (BATF, BHLHE40, and ZEB1, all known repressors) showed significant negative correlations (Supp. Fig. S8b). Since changes in predicted occupancy reflect changes in the DNase-seq data, we suspect that these repressors, upon binding to the DNA, cause the local chromatin state to become inaccessible to other factors. BATF is active in the immune system and known to interact with IRF4. Interestingly, IRF4 and BATF both had high expression in lymphoblastoid cells, yet we predicted high IRF4 occupancy and low BATF occupancy in those cells (Supp. Fig. S8). Thus, certain sets of cofactors may be utilized to up- or down-modulate the occupancy of related TFs in a cell-type- or condition-specific manner.

### TOP monitors the dynamics of TF occupancy during hormone response

Nuclear hormone receptors are TFs specifically activated in response to hormone exposure. Once activated, they bind to specific hormone response elements (HREs) where they regulate gene expression, often in conjunction with the binding of cofactors and remodeling of the chromatin structure. Glucocorticoid receptor (GR), androgen receptor (AR), and estrogen receptor (ER) are type I nuclear receptors, playing critical roles in immune response or reproductive system development, and are heavily involved in many types of cancer. To investigate TF occupancy dynamics in response to glucocorticoid, androgen, or estrogen stimulation, we predicted TF occupancy using DNase-seq data collected under each of these treatment conditions. For glucocorticoid (GC) treatment, we conducted DNase-seq experiments in A549 cells (human alveolar adenocarcinoma cell line) over 12 time points from 0 to 12 hours of GC exposure^28^. For androgen treatment, we collected DNase-seq data in LNCaP cells (human prostate adenocarcinoma cell line) over 4 time points from 0 to 12 hours following androgen induction^25^. For estrogen treatment, we used published DNase-seq data before and after estrogen induction in two kinds of cells: Ishikawa (human endometrial adenocarcinoma cell line) and T-47D (human ductal carcinoma cell line)^33^.

We identified sites with significantly differential TF occupancy before and after estrogen induction, as well as over the full time courses for GC and androgen treatment. We then ranked TFs based on the percentage of sites showing significantly increased or decreased occupancy in response to treatment. We grouped TFs with similar motifs together using RSAT clusters^34^ and present results for all significant clusters in Fig. 5 (results for individual TFs in Supp. Fig. S9). We observed different sets of TFs enriched in response to GC, androgen, and estrogen. In the list of most dynamic clusters for GC response (Fig. 5a), GR was ranked at the top—consistent with recent results showing that motif-driven GR binding is the most predictive feature of GC-inducible enhancers^28,35^—followed closely by C/EBP^28^. FOX and GATA clusters appeared next, and in both cases, while we identified more sites whose occupancy increased over the time course, we also detected a significant number that decreased.

**Fig. 5.**
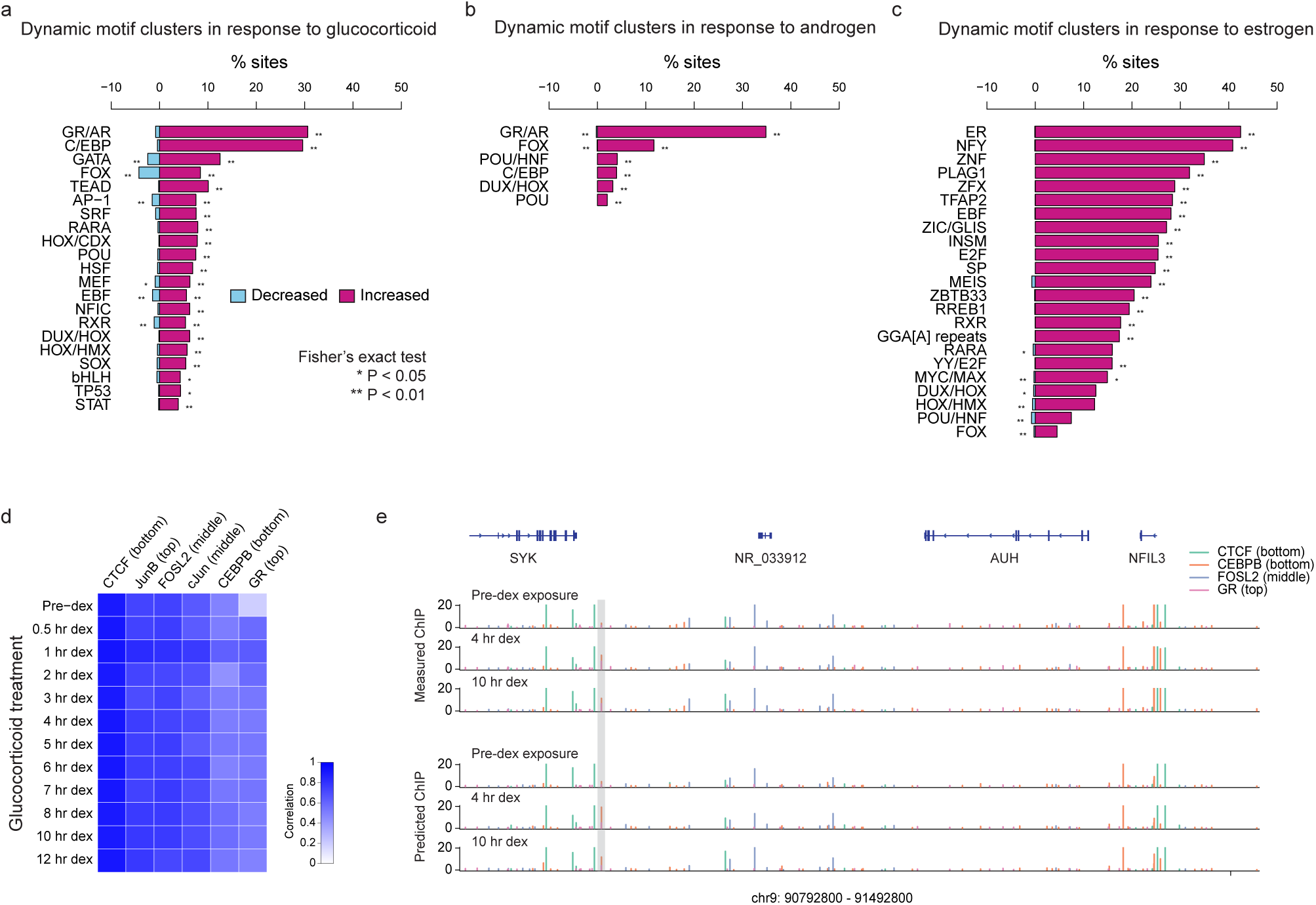
TF occupancy dynamics in response to hormone stimulation. (**a**) Motif clusters were ranked by the percentage of candidate sites whose predicted occupancy exhibited either a linear increasing or decreasing trend along the 12 time points of glucocorticoid treatment. Only significant dynamic motif clusters (*P*-value < 0.05) are listed. (**b**) Similar to (a), but along the four time points of androgen treatment. (**c**) Similar to (a) and (b), but before and after estrogen treatment. Because DNase data was collected at 12 time points during treatment with GC, at four time points with androgen, and at only two time points with estrogen, numbers are not necessarily comparable between different experiments in (a), (b), and (c). (**d**) Prediction accuracy for six TFs was evaluated afterwards using subsequently generated ChIP-seq data ^28^. Shades of blue indicate the correlation between predicted and measured occupancy for each of the six TFs at each time point. Columns (TFs) were sorted by average accuracy across the 12 time points. (**e**) Measured and predicted TF occupancy landscapes of CTCF, CEBPB, FOSL2, and GR in an example genomic region on human chromosome 9. Predicted occupancy corresponded well with measured occupancy across time, for example revealing in the highlighted region where CEBPB occupancy increased following GC treatment.

Among TFs whose occupancy was predicted to be significantly responsive to androgen treatment (Fig. 5b), AR was at the top of the list, followed by the FOX cluster. Clusters exhibited very few sites with decreasing occupancy along the time course. These observations are consistent with our previous findings that androgen induction mainly leads to an increase in chromatin accessibility, and that AR and FOXA1 are key TFs with increased occupancy^25^. The fact that the occupancy of many AR and FOXA1 sites increased gradually over the duration of the time course (Supp. Fig. S10a) highlights the importance of a quantitative perspective of TF occupancy.

In the case of estrogen induction (Fig. 5c), ER was ranked at the top, followed closely by the NFY cluster. Intriguingly, the DNase digestion profiles flanking NFYA binding sites showed striking oscillation patterns similar to those observed within nucleosomes^36^ (Supp. Fig. S11). This is consistent with previous reports that NFYA has nucleosome-like properties, and plays an important role in maintaining chromatin structure^10,37^.

That different TFs were enriched in these lists may be partly due to cell type differences, but also suggests different utilization of cofactors for GR, AR, and ER binding in response to hormone stimulation. Interestingly, we observed that PWM scores were significantly higher in sites with increased occupancy than sites of unchanged occupancy for GR, AR, and ER, but not for CEBPB, FOXA1, or NFYA (Supp. Fig. S10b), indicating that motif strength for GR, AR, and ER may play a role in prioritizing the selection of binding sites in response to hormone stimulation. This accords with recent results indicating that GR motif strength is predictive of GC-induced enhancer function^35^.

To independently validate our occupancy predictions with data not seen during training, we compared our predictions throughout the GC time course with ChIP-seq data collected in the same experiment^28^ (Fig. 5d). We computed the correlation between measured and predicted occupancies for CTCF, JunB, FOSL2, cJun, CEBPB, and GR. Across all six TFs and 12 time points, average correlation was 0.70. Over the time course, it was lowest before treatment (0.63) but otherwise consistent (between 0.68 and 0.72). Among TFs, predictions were the most accurate for CTCF (0.91)—not surprising given how predictable we observed it to be (Fig. 3)—and least for GR (0.52). Two reasons for the lower accuracy of GR are that we used a top level model because GR was not profiled as part of ENCODE, and that GR is known to have a weak DNase footprint^32^. The correlation is particularly low before treatment (time point 0), consistent with observations that many GR binding sites occur at regions of the genome that are already open prior to GC exposure^38^.

### TOP identifies genetic variants associated with predicted TF occupancy (topQTLs) and provides mechanistic interpretations for dsQTLs

A large majority of genetic variants associated with complex traits are located in non-coding genomic regions^39^, suggesting roles in transcriptional regulation. To elucidate this, it is imperative that we continue to identify genetic variants affecting TF occupancy and chromatin dynamics. To examine whether TOP is capable of sensitively distinguishing quantitatively differential TF occupancy across individuals or genetic variants, we predicted CTCF occupancy in lymphoblastoid cell lines (LCLs) from two trio studies, one from a CEU (CEPH Utah) family and one from a YRI (Yoruba from Ibadan) family^24,40^. TOP successfully identified differential CTCF occupancy between individuals across CEU and YRI families (Fig. 6a), and was sensitive enough to capture quantitative differences in CTCF occupancy between allele genotypes at allele-specific sites within CEU and YRI families (Fig. 6b).

**Fig. 6.**
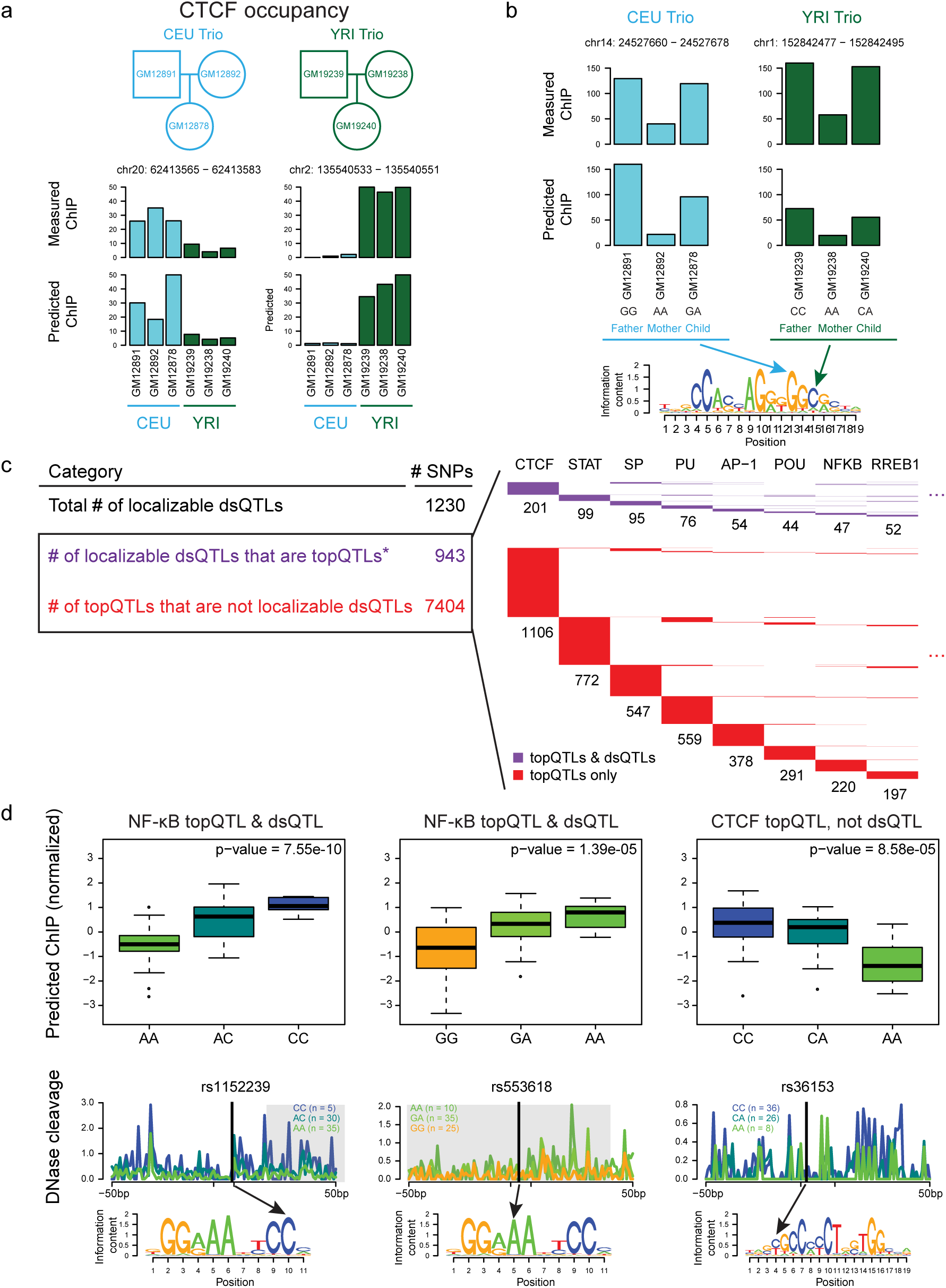
TF occupancy profile QTLs (topQTLs). (**a**) Measured and predicted CTCF occupancy at two individual-specific example loci with significantly differential CTCF occupancy between CEU and YRI families. (**b**) Measured and predicted CTCF occupancy at two allele-specific example loci with significantly differential CTCF occupancy within CEU and YRI families. (**c**) Intersections of topQTLs with ‘localizable dsQTLs’ (those within their own 100 bp windows and also within motif matches). topQTLs were defined with FDR < 10% (^⋆^ with FDR < 20%, the number of localizable dsQTLs that are also topQTLs is 1000). (Right) Largest motif clusters for topQTLs are displayed in the matrix; each row represents one topQTL which can be explained by one or more motif clusters in the columns. (**d**) Examples of topQTLs showing normalized allele-specific predicted occupancy; average DNase digestion profiles within 50 bp of the motif for each allele (significant dsQTL windows shaded in gray); and SNP locations within motifs. The CTCF topQTL overlapped a measured CTCF ChIP-seq peak in multiple LCLs, but was not identified as a dsQTL.

Encouraged by this result, we extended our predictions of genome-wide quantitative occupancy to nearly 1500 TF motifs across 70 Yoruba LCLs using TOP applied to previously published genotype and DNase-seq data^41^. With the resulting TF occupancy profiles across 70 individuals, we applied a QTL mapping strategy to identify genetic variants whose genotypes were significantly associated with changes in predicted TF occupancy, which we called ‘topQTLs’. Since genetic variants that change TF motifs often affect TF binding occupancy by changing DNA binding affinity^42^, we mapped three versions of topQTLs: within 2 kb and 200 bp *cis* testing regions around motif matches, as well as SNPs that lie strictly within those motif matches. We sought to compare our topQTLs with previously reported DNase I sensitivity quantitative trait loci (dsQTLs)^41^. We estimated the heritability and enrichment of heritability for topQTLs and dsQTLs using stratified LD score regression (S-LDSC) on publicly available GWAS summary statistics of multiple human diseases and complex traits. As shown in Supp. Fig. S12, the heritability and enrichment estimates for topQTLs are similar among different window sizes, with slightly higher enrichment for topQTLs near motif locations, consistent with our understanding of TF binding mechanisms. Heritability estimates are similar between topQTLs and dsQTLs across traits, though dsQTLs tend to show higher enrichment.

We then focused our attention on SNPs within TF motif matches because these have the highest potential for causal interpretation. We compared topQTLs within motif matches to a subset of dsQTLs that we call ‘localizable dsQTLs’, dsQTLs that fall inside the 100 bp windows with which they are linked and also lie within TF motif matches (Fig. 6c). Of the 1230 reported dsQTLs that were localizable, 943 of them were topQTLs (this number increased to 1000 when using FDR < 20%, while 1141 (93%) were associated with a significant change in predicted TF occupancy under a less stringent threshold of *P*-value < 0.05). Thus, topQTLs provide a direct mechanistic interpretation for nearly all localizable dsQTLs by revealing the identity of TFs likely to drive the observed changes in chromatin accessibility. Moreover, and importantly, we identified more than six thousand additional topQTLs that were not reported as dsQTLs. Among RSAT-clustered motifs, CTCF, STAT, SP, PU, AP-1, POU, NF-κB, and RREB1 motifs had the greatest number of topQTLs (each well over 200); most of these factors are known to be active in LCLs and critical for immune cell development^41,43^. Fig. 6d shows three sample topQTLs, one for NF-κB that is a non-localizable dsQTL, another for NF-κB that is a localizable dsQTL, and one for CTCF that is not reported as a dsQTL.

That CTCF had the largest number of topQTLs, over 1300, is noteworthy because CTCF plays a key role in chromosomal looping and commonly demarcates the boundaries of topologically associating domains (TADs)^44^. A genetic variant that disrupts a CTCF motif not only may have a significant impact on occupancy at loop anchor sites, but also could disrupt TAD boundaries. Such disruption has been demonstrated experimentally and pathologically to dysregulate the chromatin landscape and the expression of genes within the affected TAD^45^.

## Discussion

We introduce TOP to accurately predict quantitative ChIP-seq occupancy using DNase-seq data. TOP effectively learns both TF-specific and TF-generic model parameters among TFs and across cell types using a Bayesian hierarchical regression framework. TOP employs a supervised learning strategy, trained with existing TF binding specificity, DNase-seq, and ChIP-seq data, yet can accurately predict TF occupancy for new conditions, cell types, or TFs due to its hierarchical structure. In contrast to traditional ways of analyzing ChIP-seq data through peak-calling to label genomic regions as bound or unbound, TOP adopts a quantitative perspective, allowing us to predict the level of TF occupancy along a continuum. This opens up a new way to investigate quantitative changes in TF occupancy across cell types, treatment conditions, and developmental time courses. TOP is general in that it can predict occupancy for any sequence-specific TF of interest with any new DNase-seq data in any cell type or condition without requiring a new ChIP-seq experiment. TOP’s ability to use time-course DNase data over 12 hours of GC treatment served as a cost-effective strategy to study the temporal dynamics of TF occupancy. By doing one DNase-seq experiment at each time point, we obtained occupancy predictions for 1500 TF motifs, allowing us to screen for TFs showing significant changes in occupancy. For example, TOP results suggest a significant role for FOX and GATA factors in GC-induced transcriptional response (Fig. 5a). Although developed for DNase-seq data, TOP can easily be extended to ATAC-seq data, and though it was trained on human data, it is equally applicable in other organisms. As an example demonstration, we showed that it can successfully predict quantitative Reb1 occupancy across the yeast genome (Supp. Fig. S13). As a resource for the community, we provide genome browser tracks of predicted occupancy for nearly 1500 motifs across 178 cell types, throughout a 12-hour time course of GC exposure, and across 70 LCLs. The occupancy map of TF × cell-type combinations alone expands the total output of ENCODE TF ChIP-seq efforts over 200-fold.

Recently, methods have emerged for the imputation of missing epigenomic data (histone modifications, chromatin accessibility, etc.) using the many other types of available data generated by the ENCODE consortium^46–48^. Our approach shares a similar goal with these imputation methods in trying to predict unmeasured data using models trained on existing datasets across multiple cell types. However, we note some major distinctions between our approach and these recent imputation strategies. First, our approach requires only DNase-seq (or ATAC-seq) and TF ChIP-seq data for training, and requires only DNase-seq (or ATAC-seq) data to make predictions. In contrast, existing imputation methods often require a large variety of existing assays (DNase-seq, RNA-seq, histone modification ChIP-seq, etc.), which may not be readily available, especially in studies to profile new cell types or treatment conditions. Second, our strategy predicts TF occupancy only at candidate binding sites (based on low stringency motif matches), whereas existing imputation approaches attempt to impute a TF’s ChIP-seq signal across the entire genome, devoting statistical power and computational effort to genomic locations where a given TF is not likely to bind, which is the vast majority. Third, TOP uses a Bayesian hierarchical regression framework to model DNase digestion features, which allows for easier interpretation than more complex methods, especially those involving deep neural networks or tensor factorization. Last but not least, our hierarchical model is able to predict the binding of TFs that have never been measured by ChIP-seq, a significant advantage over imputation methods that require ChIP-seq training data for TFs of interest.

The fact that TOP predicts TF occupancy only at candidate binding sites is also a limitation because it has been observed that for many TFs, a large number of their ChIP-seq peaks do not have motif matches^42^. On the other hand, this does have some benefits, since distinguishing direct from indirect binding can be difficult using ChIP assays. By focusing only on motif matches, our results can be viewed as predictions of TF occupancy that are explainable by direct binding. Indeed, TOP could be used to suggest direct-binding TFs that may be mediating the indirect binding of other TFs in ChIP experiments. As a last observation, since motif quality directly affects which genomic locations we select as candidate binding sites, and since PWM scores also factor into TOP predictions, better TF binding affinity models are important, and should improve TOP’s predictions in the future.

Our approach can be viewed as complementary to ChIP-based exploration of TF occupancy. Instead of doing one ChIP-seq experiment for every TF in a particular cell type or condition, TOP needs only one DNase-seq experiment to predict the genome-wide occupancy of many TFs. TOP can therefore be used to screen and identify TFs showing significant changes in occupancy, enabling the prioritization of future ChIP experiments for a small number of key TFs. The modeling strategy we present here offers a foundational and cost-effective approach for profiling the quantitative occupancy of myriad TFs across diverse cell types, dynamic conditions, and genetic variants.

## Methods

### Candidate binding site selection

We defined candidate TF binding sites by PWM scanning across the genome using FIMO^29^. When training or applying our model, we included as candidate sites all motif matches with *P*-value < 10^−5^. Similar to CENTIPEDE ^5^ and MILLIPEDE ^6^, we filtered out candidate sites if more than 10% of the nucleotides in the surrounding window (100 bp flanking each side of the motif) were unmappable.

When training the regression model, if the training TF had more than one motif, we manually selected one based on which was the most representative motif for that TF in the Factorbook database^49^. After training, we used the model parameters estimated at various levels of the TOP hierarchy to make occupancy predictions for 1496 motifs, including both JASPAR core motifs (2014 version) and those used by Sherwood and colleagues^10^. The motifs selected for training the model, along with the full list of all motifs used for prediction in this paper, are provided in Supp. Tables S2 and S3.

### Normalization and data preprocessing

To account for differences in sequencing depth across experiments in different cell types or conditions, DNase-seq and ChIP-seq data were normalized by library size (scaled to library sizes of 100 million mapped reads for DNase-seq and 10 million mapped reads for ChIP-seq data). This simple library size normalization is flexible for downstream analysis. We considered other types of normalization methods, including quantile normalization, trimmed mean of M-values (TMM), etc. However, these methods assume different experiments will have the same distribution of reads across peaks (or a subset of common peaks) among all the experiments, which is too strong an assumption in our case—especially, for example, when comparing hormone receptor binding before and after hormone induction—and leads to a high number of false negatives in the GR, AR, and ER analyses.

### DNase feature extraction using binning vs. wavelet coefficients

We systematically evaluated different features of the cleavage events (‘cuts’) arising from DNase digestion, in an attempt to avoid overfitting^11^ and any possible influence of DNase digestion bias. First, we tried to extract multi-resolution features of DNase digestion data using wavelet multi-resolution decomposition. Wavelet methods provide a natural approach to extract the multi-resolution information contained in both DNase cut magnitude and detail profiles. Here, we decomposed DNase-seq data using Haar wavelets with the `wavethresh` package in R. The detail signals were extracted at different resolution levels through the mother wavelet coefficients while the scales of cuts at different resolution levels were represented by the father wavelet coefficients. We started with windows of size 128 bp around the motif center, but later focused on 64 bp windows around the motif centers, because that was where the majority of the largest mother wavelet coefficients were located. Then we fit regression models with mother wavelet coefficients and log-transformed father wavelet coefficients at multiple resolution levels as predictors, together with PWM score, and conducted variable selection with LASSO. Interestingly, variable selection results suggested the scale of DNase cuts (represented by the father wavelet coefficients) was the most significant feature for predicting TF occupancy (Supp. Fig. S1), consistent with previous findings^6,8,50^. In contrast, very few spiky DNase signals (represented by the mother wavelet coefficients) were selected. Worse, some of the fine details in the DNase signal in the motif region might arise from sequence-specific DNase digestion bias^6,8,30,50^.

Based on these empirical observations of DNase digestion profiles around motifs, we simplified the process of DNase feature extraction by using a more flexible binning scheme in place of the rigid dyadic splitting of the wavelet framework. In previous work, we developed a model called MILLIPEDE that divides the motif region and its flanking regions upstream and down-stream into various distinct bins^6^ (Fig. S2a). Following the binning scheme of MILLIPEDE, we compared different binning models from the most complicated M12 model to the simplest M1 model, and evaluated their performance in comparison to an optimally-selected wavelet model. Fig. S2b shows the prediction performance of all these models for four TFs in K562 cells using 5-fold cross-validation. In summary, different binning models led to roughly similar prediction performances and were generally comparable to a model using optimally-selected wavelet features. In agreement with our earlier results^6^, M5 binning—which effectively summarizes the number of DNase cleavage events in the motif region, nearby flanking regions, and distal flanking regions on both sides of the motif—is complex enough to capture the DNase digestion features sufficient to predict TF occupancy at high accuracy, but is also simple enough to fit into the Bayesian hierarchical regression framework and still yield easily interpretable TF-specific and TF-generic signatures. Fig. 2 shows the prediction performance of the TOP model using M5 binning on the training data. We observed very close agreement in prediction performance using training data vs. test results from cross-validation for the TFs listed in Fig. S2b.

### Bayesian hierarchical regression model

We designed the hierarchical model to have three levels, with cell types nested within TF branches. (In principle, we could expand the hierarchical model to have an additional branch with parameters for each cell type, i.e., a cell-type–specific but TF-generic model. However, we expect a TF to have similar model parameters in different cell types. Also, in most of the ENCODE tier 3 cell types, very few TFs have been profiled with ChIP-seq, so we would likely have insufficient data to estimate cell-type–specific parameters for most cell types.)

ChIP-seq count data are typically fit using a negative binomial distribution, which uses an extra parameter to model the overdispersion in ChIP-seq data better than a Poisson distribution. However, we found a simpler Gaussian linear model on asinh-transformed ChIP-seq data to be a better choice for fitting our Bayesian hierarchical model (asinh transformation is similar to log transformation but handles zero values more gracefully; we used it successfully in our recent NucID model^36^). This choice has the added benefit of applying to non-integer data, which arise whenever we average counts over replicate experiments or conduct data normalization. We compared the prediction accuracy of our Gaussian linear model on asinh-transformed data against the alternative of negative binomial regression on integer-rounded data, and observed very close agreement. We ultimately decided to use the Gaussian distribution because it has a nice conjugacy property, allowing posteriors to be estimated through Gibbs sampling, thereby providing a computational advantage over a negative binomial distribution.

The basic regression model for modeling the asinh-transformed ChIP-seq occupancy *y*_*t, c, i*_ that is observed when TF *t* occupies its candidate binding site *i* in cell type *c* can be briefly summarized as:

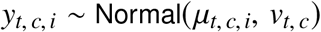

Where

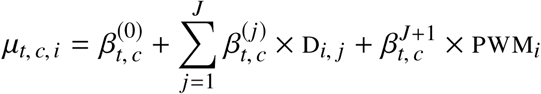

The D_*i, j*_ variable represents DNase feature *j* for site *i*, while *J* is the number of DNase features in the model (in our final model, we use M5 binning so *J* = 5).

The Bayesian hierarchical model is specified as follows:

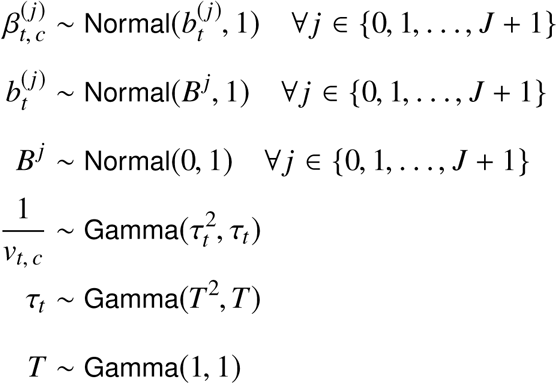

We tested whether allowing the variances of the various beta parameters to be learned from the data using an inverse gamma hyperprior, but the results were essentially unchanged. We used the consensus Monte Carlo algorithm^51^, a parallel technique to reduce the running time of the Gibbs sampler while maintaining predictive performance. Briefly, we split all data randomly into ten equal parts. Gibbs samplers were run on each part separately in parallel for 10^6^ iterations. Posterior samples from the first 8 × 10^5^ iterations were discarded for burn-in. Each of the remaining 2 × 10^5^ posterior samples from the ten Gibbs samplers were averaged to get the final posterior samples for each model’s parameters.

### Comparison of prediction accuracy with existing methods

We compared TOP with four existing methods. CENTIPEDE and msCentipede predict TF binding probabilities using an unsupervised generative framework to model the DNase digestion profiles around candidate sites (motif matches) without ChIP-seq training data. msCentipede improves on CENTIPEDE by using a multi-scale model framework to better model heterogeneity across sites and replicates. We ran CENTIPEDE and msCentipede on DNase cuts data in each TF-cell type combination under default parameter settings. CENTIPEDE was run on DNase data after pooling the replicate samples. msCentipede was run on individual DNase replicates to better capture heterogeneity (its authors demonstrated how beneficial replicates are to msCentipede accuracy). Because the CENTIPEDE paper showed a substantial correlation between its TF binding predictions (posterior log odds) and ChIP-seq read counts (sqrt transformed), we computed Pearson correlations between measured ChIP-seq counts (library size normalized and sqrt transformed) and posterior log odds from CENTIPEDE and msCentipede, as well as to the quantitative predictions made by TOP models at each level of the hierarchy. In contrast to CENTIPEDE and msCentipede, MILLIPEDE^6^ adopts a supervised learning strategy using a logistic regression framework with binary ChIP-seq peaks as training labels and TF-generic binning to extract DNase digestion features (similar to TOP). BinDNase^12^ is a later method that is very similar to MILLIPEDE, but allows each TF to have its own DNase binning scheme. We ran MILLIPEDE (with M5 binning) and BinDNase on DNase cuts data in each TF-cell type combination under default parameter settings. We did 5-fold cross-validation to evaluate the prediction performance by splitting the chromosomes into five groups of similar sizes. To predict TF binding for each chromosome group (test set), we trained the models using the remaining four chromosome groups (training set). As with CENTIPEDE and msCentipede, we computed the Pearson correlations between the log odds of TF binding probabilities and sqrt transformed ChIP-seq read counts.

We did not include PIQ^10^ in our comparison, because msCentipede has already been shown to significantly outperform PIQ when it has access to DNase replicates^11^. GERV^13^ is a statistical method that learns a k-mer based model to predict TF binding using ChIP-seq and DNase-seq data and scores genetic variants by quantifying the changes of predicted ChIP-seq reads between the reference and alternative allele. Like TOP, it tries to predict quantitative TF occupancy, but its main goal is to score genetic variants that affect TF binding, and it treats DNase signals as a binary feature (open vs. closed), which would not be effective in capturing quantitative changes in DNase signals across dynamic conditions. Also, as a k-mer based method, it does not adopt the motif-centric framework that we and the other methods do. For these reasons, we did not include GERV in our comparison.

### Differential occupancy comparison across cell types

We used the `edgeR` package^52^ to identify sites with significantly differential occupancy across cell types. For each TF at each candidate binding site, we tested the cell-type effect by contrasting the predicted occupancy in each cell type (using DNase replicate samples) against the cell-type mean. Sites with predicted occupancy less than 1 read per million were filtered out from the test, and then sites with a significant cell-type effect (FDR < 10%) were selected.

When comparing predicted occupancy across cell types, potential influences from copy number variation (CNV) could lead to false positives. However, since our method predicts TF occupancy using DNase data, and since CNV affects both DNase-seq and ChIP-seq counts in a consistent manner (CNV would lead to higher occupancy in both measured and predicted ChIP-seq in a higher copy number region), our predictions should still agree with measured occupancy. To deal with CNV influences while comparing across cell types, instead of directly correcting CNV on both DNase-seq and ChIP-seq data within the regression model, it is easier to do CNV adjustment as a post-processing procedure on the predicted occupancy using input ChIP-seq data. However, since not all these cell types have input ChIP-seq data available, we did not perform CNV corrections in this study (input correction could be performed in those cell types for which input ChIP-seq data are available).

### DNase-seq data across hormone treatment conditions

DNase-seq data from LNCaP cells exposed to androgen were collected in our labs. Data from before induction (time point 0) and after 12 hours were already previously published^25^ and are available from the Gene Expression Omnibus (GEO) repository under accession GSE34780. DNase-seq data from the 45 minute and 4 hour treatments, along with more samples from before induction, were generated for this study and is available from the GEO repository under accession GSE157473.

LNCaP cells were obtained from ATCC. Cells were maintained using the protocol described at http://genome.ucsc.edu/ENCODE/protocols/cell/human/LNCaP_Crawford_protocol.pdf. Prior to stimulation with either androgen (R1881, methyltrienolone) or vehicle (ethanol) for varying time durations, cells were grown in RPMI-1640 medium with 10% charcoal:dextran stripped medium for 60 hours. Androgen was added to the culture medium for a final concentration of 1 nM in all experiments. Isolation of total DNA, cleavage with DNase I (henceforth, DNase), and subsequent preparation of sequencing libraries were carried out as previously described (Song and Crawford, Cold Spring Harbor Protocol 2010). Replicates from 12 hours of androgen exposure were previously sequenced on the Illumina GAIIx platform, whereas replicates from the 45 minute and 4 hour time points were sequenced for this study on the Illumina HiSeq2000 platform. Sequenced reads were aligned to the genome and further processed as previously described^25,53,54^.

DNase-seq data from A549 cells exposed to the glucocorticoid hormone dexamethasone were collected in our labs. Detailed methods are provided in our paper^28^.

DNase-seq data from Ishikawa and T-47D cells before and after estrogen exposure were collected by others and previously published^33^; we downloaded their published data.

### Differential occupancy comparison across hormone treatment conditions

In the androgen treatment analysis, we combined DNase-seq data from an earlier study^25^ with three replicates of uninduced samples and two replicates of 12 hour androgen induced samples (using the Illumina GAIIx sequencing platform), and DNase-seq data generated in this study with two replicates of uninduced samples, two replicates of 45 minute induced samples, and two replicates of 4 hour induced samples (using the Illumina HiSeq2000 sequencing platform). AR ChIP-seq data collected in an earlier study with 4 hour androgen induction in LNCaP cells^55^ matched with our DNase-seq data of 4 hour androgen induction were included in the training dataset for AR in the hierarchical model. For each TF, we used `edgeR` to test linear, quadratic, and cubic trends of TF occupancy changes over the time course of uninduced, 45 minute, 4 hour, and 12 hour induced conditions, adjusting for the batch effect from different sequencing platforms (GAIIx vs. HiSeq sequencing). Sites with predicted occupancy less than 10 were filtered out from the test, and then sites with significant linear, quadratic, or cubic trend of TF occupancy over the time course (FDR < 10%) were selected. Very few sites were found to have a significant quadratic or cubic trend, so we focused on sites with a significant linear trend.

In the estrogen treatment analysis, we used previously published DNase-seq and ChIP-seq data generated in Ishikawa (endometrial cancer cell line; previously mislabeled as ECC-1) and T-47D (breast cancer cell line) cells before and after estrogen induction^33^. ER ChIP-seq data from estrogen induced conditions were matched with the corresponding DNase data and included in the training dataset for ER in the hierarchical model. Occupancy predictions were made for each TF using its middle level parameters in DNase-seq replicate samples in Ishikawa and T-47D, before and after estrogen stimulation. For each TF, we used `edgeR` to test for differential occupancy, where we considered both cell-type effect (Ishikawa vs. T-47D) and treatment effect (estrogen induced vs. uninduced). Sites with predicted occupancy less than 10 were filtered out from the test, and then sites with treatment effect significantly higher or lower than zero (FDR < 10%) were selected.

In the glucocorticoid (GC) treatment analysis, we used DNase-seq data collected in our labs from A549 cells (human alveolar adenocarcinoma cell line) over 12 time points from 0 to 12 hours following exposure to the glucocorticoid hormone dexamethasone^28^. For each TF, we used `edgeR` to test linear, quadratic, and cubic trends of TF occupancy changes over the 12 time points of GC treatment. Sites with predicted occupancy less than 10 were filtered out from the test, and then sites with significant linear, quadratic, or cubic trend of TF occupancy over the time course (FDR < 10%) were selected. Very few sites were found to have a significant quadratic or cubic trend, so we focused on sites with a significant linear trend.

After selecting sites with significant differential occupancy, we ranked TFs based on the percentage of sites showing significantly increased or decreased occupancy in response to treatment. TFs with similar motifs were grouped together using RSAT clusters^34^ to simplify downstream interpretation and visualization.

### topQTL mapping

We predicted genome wide TF occupancy for 1496 motifs using previously published genotype information and DNase data generated from LCLs from 70 individuals^41^. For each motif, we focused on those motif matches that had a SNP inside. When making predictions across the 70 LCLs using both PWM scores and DNase data, we fixed the PWM scores for candidate sites to be the average of the PWM scores calculated from the two homozygous genotypes for that SNP, in order to avoid using PWM scores twice: in both occupancy predictions and QTL association testing. We mapped topQTLs by testing the associations between genotypes and predicted TF occupancy across the 70 individuals using a linear model (with R package `MatrixEQTL`). For each TF motif, we selected the 10% of candidate sites with the highest predicted occupancy for QTL mapping and downstream analysis (we tested top 10%, 20%, …, 100% sites, and found the top 10% sites tended to maximize the number of QTLs detected after multiple testing correction). To facilitate comparison with dsQTLs, we followed the same data processing procedures as described by the authors^41^, including z-score standardization, quantile normalization, and regressing out 4 PCs to remove unidentified confounders. Following Degner *et al*.^41^, we corrected the effect of GC content by first partitioning candidate windows (motif matches plus 100 bp flanking windows on both sides) into bins according to their GC content and then normalizing each sample by subtracting each bin median from all the windows belonging to the partition of the corresponding bin. We mapped three versions of *cis* topQTLs by testing SNPs in 200 bp and 2 kb windows around candidate sites, as well as SNPs within motif matches. For each of the TF motifs, genetic variants with significant associations to predicted TF occupancy (FDR < 10%) were identified as topQTLs for that TF motif, and were the basis of all subsequent analysis in Fig. 6 of the manuscript. TFs with similar motifs were grouped together using RSAT clusters^34^ to simplify downstream interpretation and visualization.

### Heritability and enrichment analysis of GWAS summary statistics using S-LDSC

We partitioned the heritability of complex traits and estimated heritability enrichment of topQTLs and dsQTLs using S-LDSC^56^. S-LDSC partitions the heritability of genomic annotations using GWAS summary statistics and estimates the enrichment as a ratio of the proportion of heritability explained by an annotation divided by the proportion of SNPs in that annotation. We constructed binary annotations containing lead SNPs of topQTLs (using 200 bp and 2 kb *cis*-testing regions around motif matches, as well as SNPs within motifs) and lead SNPs of dsQTLs downloaded from Degner et al.^41^ (using 2 kb and 40 kb *cis*-testing regions around the 5% of 100 bp DNase windows with the highest DNase I sensitivity). An FDR threshold of 10% was used for both topQTLs and dsQTLs. We applied S-LDSC to our QTL-based annotations using separate models for each QTL annotation. In our S-LDSC analysis, we adjusted for various baseline annotations of SNPs using a baselineLD model^57^, including gene annotations (coding, UTRs, intron, promoter), minor allele frequency, and LD-related annotations. We did not include functional annotations such as enhancer marks in our baseline model, since these annotations are likely correlated with the QTL features of interest, and including them may bias our estimates. The GWAS traits and corresponding references are listed in Supp. Table S4.

## Data availability

DNase-seq data in LNCaP cells before and after 45 minutes and 4 hours of 1 nM R1881 treatment have been deposited in the Gene Expression Omnibus (GEO) repository under accession GSE157473. Pre-computed genome-wide tracks of quantitative TF occupancy will be made available upon publication at http://www.cs.duke.edu/~amink/software/.

## Code availability

TOP is implemented in R, and all code will be made available on GitHub upon publication. Pre-trained TOP model parameters, and links to all code resources will also be made available from a single location upon publication at http://www.cs.duke.edu/~amink/software/.

## Acknowledgements

The authors would like to particularly thank David MacAlpine, Raluca Gordaân, Galip Guürkan Yardimci, Jason Belsky, Ian McDowell, Chris Vockley, Tony D’Ippolito, and the rest of the Duke GGR team for helpful comments during the development of this work or in response to drafts of the manuscript. This work was funded in part by NIH grants U01-HG007900 and R01-GM118551.

## Author Contributions

K.L., G.E.C., and A.J.H. designed the study. K.L., J.Z., L.M., G.E.C., and A.J.H. conducted and supervised analyses. A.S., L.K.H, A.K.T., L.S., T.E.R., and G.E.C. conducted and supervised experiments. K.L., J.Z., and A.J.H. wrote the paper.

## Competing interests

The authors declare no competing interests.

## Notes

### Competing Interest Statement

The authors have declared no competing interest.

## References

[1] Thurman, R. E. et al. The accessible chromatin landscape of the human genome. Nature 489, 75–82 (2012).

[2] Neph, S. et al. An expansive human regulatory lexicon encoded in transcription factor footprints. Nature 489, 83–90 (2012).

[3] Buenrostro, J. D., Wu, B., Chang, H. Y. & Greenleaf, W. J. ATAC-seq: A method for assaying chromatin accessibility Genome-Wide. Curr. Protoc. Mol. Biol. 109, 21.29.1–21.29.9 (2015).

[4] Klemm, S. L., Shipony, Z. & Greenleaf, W. J. Chromatin accessibility and the regulatory epigenome. Nat. Rev. Genet. 20, 207–220 (2019).

[5] Pique-Regi, R. et al. Accurate inference of transcription factor binding from DNA sequence and chromatin accessibility data. Genome Res. 21, 447–455 (2011).

[6] Luo, K. & Hartemink, A. J. Using DNase digestion data to accurately identify transcription factor binding sites. In Pac. Symp. Biocomputing, 80–91 (World Scientific, Hackensack, NJ, 2013).

[7] He, H. H. et al. Differential DNase I hypersensitivity reveals factor-dependent chromatin dynamics. Genome Res. 22, 1015–1025 (2012).

[8] He, H. H. et al. Refined DNase-seq protocol and data analysis reveals intrinsic bias in transcription factor footprint identification. Nat. Methods 11, 73–78 (2014).

[9] Zhong, J., Wasson, T. & Hartemink, A. J. Learning protein-DNA interaction landscapes by integrating experimental data through computational models. Bioinformatics 30, 2868–2874 (2014).

[10] Sherwood, R. I. et al. Discovery of directional and nondirectional pioneer transcription factors by modeling DNase profile magnitude and shape. Nat. Biotechnol. 32, 171–178 (2014).

[11] Raj, A., Shim, H., Gilad, Y., Pritchard, J. K. & Stephens, M. msCentipede: Modeling heterogeneity across genomic sites and replicates improves accuracy in the inference of transcription factor binding. PLoS ONE 10, e0138030 (2015).

[12] Kähärä, J. & Lähdesmäki, H. BinDNase: A discriminatory approach for transcription factor binding prediction using DNase I hypersensitivity data. Bioinformatics 31, 2852–2859 (2015).

[13] Zeng, H., Hashimoto, T., Kang, D. D. & Gifford, D. K. GERV: A statistical method for generative evaluation of regulatory variants for transcription factor binding. Bioinformatics 32, 490–496 (2016).

[14] Li, H., Quang, D. & Guan, Y. Anchor: Trans-cell type prediction of transcription factor binding sites. Genome Res. (2018).

[15] Keilwagen, J., Posch, S. & Grau, J. Accurate prediction of cell type-specific transcription factor binding. Genome Biol. 20, 505 (2019).

[16] Quang, D. & Xie, X. FactorNet: A deep learning framework for predicting cell type specific transcription factor binding from nucleotide-resolution sequential data. Methods 166, 40–47 (2019).

[17] Schreiber, J., Bilmes, J. & Noble, W. S. Completing the ENCODE3 compendium yields accurate imputations across a variety of assays and human biosamples. Genome Biol. 21, 82 (2020).

[18] Hoffman, M. M. et al. Unsupervised pattern discovery in human chromatin structure through genomic segmentation. Nat. Methods 9, 473–476 (2012).

[19] Narlikar, L., Gordân, R. & Hartemink, A. J. Nucleosome occupancy information improves de novo motif discovery. International Conference on Research in Computational Molecular Biology (RECOMB 2007) 107–121 (2007).

[20] Gordân, R. & Hartemink, A. J. Using DNA duplex stability information for transcription factor binding site discovery. In Pac. Symp. Biocomputing, 453–464 (World Scientific, Hackensack, NJ, 2008).

[21] Wasson, T. & Hartemink, A. J. An ensemble model of competitive multi-factor binding of the genome. Genome Res. 19, 2101–2112 (2009).

[22] Li, X.-Y. et al. The role of chromatin accessibility in directing the widespread, overlapping patterns of Drosophila transcription factor binding. Genome Biol. 12, R34 (2011).

[23] Lickwar, C. R., Mueller, F., Hanlon, S. E., McNally, J. G. & Lieb, J. D. Genome-wide protein-DNA binding dynamics suggest a molecular clutch for transcription factor function. Nature 484, 251–255 (2012).

[24] McDaniell, R. et al. Heritable individual-specific and allele-specific chromatin signatures in humans. Science 328, 235–239 (2010).

[25] Tewari, A. K. et al. Chromatin accessibility reveals insights into androgen receptor activation and transcriptional specificity. Genome Biol. 13, R88 (2012).

[26] Gertz, J., Siggia, E. D. & Cohen, B. A. Analysis of combinatorial cis-regulation in synthetic and genomic promoters. Nature 457, 215–218 (2009).

[27] Segal, E. & Widom, J. From DNA sequence to transcriptional behaviour: A quantitative approach. Nat. Rev. Genet. 10, 443–456 (2009).

[28] McDowell, I. C. et al. Glucocorticoid receptor recruits to enhancers and drives activation by motif-directed binding. Genome Res. 28, 1272–1284 (2018).

[29] Grant, C. E., Bailey, T. L. & Noble, W. S. FIMO: Scanning for occurrences of a given motif. Bioinformatics 27, 1017–1018 (2011).

[30] Sung, M.-H., Guertin, M. J., Baek, S. & Hager, G. L. DNase footprint signatures are dictated by factor dynamics and DNA sequence. Mol. Cell 56, 275–285 (2014).

[31] Voss, T. C. et al. Dynamic exchange at regulatory elements during chromatin remodeling underlies assisted loading mechanism. Cell 146, 544–554 (2011).

[32] Goldstein, I. et al. Transcription factor assisted loading and enhancer dynamics dictate the hepatic fasting response. Genome Res. 27, 427–439 (2017).

[33] Gertz, J. et al. Distinct properties of cell-type-specific and shared transcription factor binding sites. Mol. Cell 52, 25–36 (2013).

[34] Castro-Mondragon, J., Jaeger, S., Thieffry, D., Thomas-Chollier, M. & van Helden, J. RSAT matrix-clustering: Dynamic exploration and redundancy reduction of transcription factor binding motif collections. bioRxiv 065565 (2016).

[35] Vockley, C. M. et al. Direct GR binding sites potentiate clusters of TF binding across the human genome. Cell 166, 1269–1281.e19 (2016).

[36] Zhong, J. et al. Mapping nucleosome positions using DNase-seq. Genome Res. 26, 351–364 (2016).

[37] Nardini, M. et al. Sequence-specific transcription factor NF-Y displays histone-like DNA binding and H2B-like ubiquitination. Cell 152, 132–143 (2013).

[38] Reddy, T. E., Gertz, J., Crawford, G. E., Garabedian, M. J. & Myers, R. M. The hypersensitive glucocorticoid response specifically regulates Period 1 and expression of circadian genes. Mol. Cell. Biol. 32, 3756–3767 (2012).

[39] Hindorff, L. A. et al. Potential etiologic and functional implications of genome-wide association loci for human diseases and traits. Proc. Natl. Acad. Sci. USA 106, 9362–9367 (2009).

[40] 1000 Genomes Project Consortium et al. A map of human genome variation from population-scale sequencing. Nature 467, 1061–1073 (2010).

[41] Degner, J. F. et al. DNaseI sensitivity QTLs are a major determinant of human expression variation. Nature 482, 390–394 (2012).

[42] Deplancke, B., Alpern, D. & Gardeux, V. The genetics of transcription factor DNA binding variation. Cell 166, 538–554 (2016).

[43] Tehranchi, A. K. et al. Pooled ChIP-seq links variation in transcription factor binding to complex disease risk. Cell 165, 730–741 (2016).

[44] Rao, S. S. P. et al. A 3D map of the human genome at kilobase resolution reveals principles of chromatin looping. Cell 159, 1665–1680 (2014).

[45] Lupiáñez, D. G., Spielmann, M. & Mundlos, S. Breaking TADs: How alterations of chromatin domains result in disease. Trends in Genetics 32, 225–237 (2016).

[46] Ernst, J. & Kellis, M. Large-scale imputation of epigenomic datasets for systematic annotation of diverse human tissues. Nat. Biotechnol. 33, 364–376 (2015).

[47] Durham, T. J., Libbrecht, M. W., Howbert, J. J., Bilmes, J. & Noble, W. S. PREDICTD PaRallel Epigenomics Data Imputation with Cloud-based Tensor Decomposition. Nat. Commun. 9, 1–15 (2018).

[48] Schreiber, J., Durham, T., Bilmes, J. & Noble, W. S. Avocado: a multi-scale deep tensor factorization method learns a latent representation of the human epigenome. Genome Biol. 21, 81–18 (2020).

[49] Wang, J. et al. Sequence features and chromatin structure around the genomic regions bound by 119 human transcription factors. Genome Res. 22, 1798–1812 (2012).

[50] Cuellar-Partida, G. et al. Epigenetic priors for identifying active transcription factor binding sites. Bioinformatics 28, 56–62 (2012).

[51] Scott, S. L. et al. Bayes and big data: The consensus Monte Carlo algorithm. Intl. J. Manage. Sci. Engin. Manage. 11, 78–88 (2016).

[52] Robinson, M. D., McCarthy, D. J. & Smyth, G. K. edgeR: A Bioconductor package for differential expression analysis of digital gene expression data. Bioinformatics 26, 139–140 (2010).

[53] Boyle, A. P., Guinney, J., Crawford, G. E. & Furey, T. S. F-Seq: A feature density estimator for high-throughput sequence tags. Bioinformatics 24, 2537–2538 (2008).

[54] Yardimci, G. G., Frank, C. L., Crawford, G. E. & Ohler, U. Explicit DNase sequence bias modeling enables high-resolution transcription factor footprint detection. Nucleic Acids Res. 42, 11865–11878 (2014).

[55] Massie, C. E. et al. The androgen receptor fuels prostate cancer by regulating central metabolism and biosynthesis. EMBO J. 30, 2719–2733 (2011).

[56] Finucane, H. K. et al. Partitioning heritability by functional annotation using genome-wide association summary statistics. Nat. Genet. 47, 1228–1235 (2015).

[57] Gazal, S. et al. Linkage disequilibrium-dependent architecture of human complex traits shows action of negative selection. Nat. Genet. 49, 1421–1427 (2017).

[58] Sheffield, N. C. et al. Patterns of regulatory activity across diverse human cell types predict tissue identity, transcription factor binding, and long-range interactions. Genome Res. 23, 777–788 (2013).

[59] Bar-Joseph, Z., Gifford, D. K. & Jaakkola, T. S. Fast optimal leaf ordering for hierarchical clustering. Bioinformatics 17 Suppl 1, S22–9 (2001).

