## Supplementary Materials for "Quantitative occupancy of myriad transcription factors from one DNase experiment enables efficient comparisons across conditions"

### Supplementary tables

#### Supplementary Table S1. Different modeling frameworks for predicting TF binding using DNase-seq data.

| | CENTIPEDE (2012) | MILLIPEDE (2013) | PIQ (2014) | BinDNase (2015) | msCentipede (2015) | TOP (this study) | GERV (2015) | Avocado (2020) |
|---|---|---|---|---|---|---|---|---|
| **Has access to TF motifs as input?** | Yes | Yes | Yes | Yes | Yes | Yes | No, uses k-mers instead | No |
| **Genome-scale data used as input for TF prediction** | One DNase-seq dataset | One DNase-seq dataset (ChIP-seq data used in training, but not used in prediction) | One DNase-seq dataset | One DNase-seq dataset (ChIP-seq data used in training, but not used in prediction) | One DNase-seq dataset (but higher accuracy with replicates) | One DNase-seq dataset (ChIP-seq data used in training, but not used in prediction) | Sequence variants (DNase- and ChIP-seq data used in training, but not used in prediction) | Trained on available ENCODE datasets including DNase-seq, ATAC-seq, RNA-seq, and ChIP-seq both for TFs and for histone modifications. Transfer learning could be used for predicting new samples. |
| **Motif/site-centric vs. genome-wide** | Motif/site-centric | Motif/site-centric | Motifs modeled jointly | Motif/site-centric | Motif/site-centric | Motif/site-centric | Genome-wide | Genome-wide |
| **Supervised vs. unsupervised** | Unsupervised | Supervised | Unsupervised | Supervised | Unsupervised | Supervised | Supervised | Supervised |
| **Independent model for each TF vs. joint model for TFs and/or cell types** | Independent model for each TF | Independent model for each TF | Joint model for all TFs in each cell type | Independent model for each TF | Independent model for each TF | Joint model for all TFs in all cell types using a Bayesian hierarchical framework | Independent model for each TF | Joint model for all types of observations in all cell types |
| **How DNase data are used in the model** | Nucleotide-resolution DNase digestion profile around motifs | Binned DNase digestion profile around motifs (same binning scheme for all TFs) | Nucleotide-resolution GP-smoothed DNase digestion profile around motifs | Binned DNase digestion profile around motifs (different binning scheme for each TF) | Multiscale nucleotide-resolution DNase digestion profile around motifs | Binned DNase digestion profile around motifs (same binning scheme for all TFs) | Nucleotide-resolution binary (0/1) indicator variable denoting presence of a DNase read across genome | Multiscale binned DNase digestion profile across genome (at 25bp, 250bp, and 5000bp resolutions) |
| **Output type and interpretation** | TF binding probability | TF binding probability | TF binding probability | TF binding probability | TF binding probability | Quantitative ChIP-seq signal expressed as counts (in window ±100bp around motif) | Quantitative ChIP-seq signal expressed as counts (in window ±200bp around k-mer) | Quantitative ChIP-seq signal expressed as signal *p* value |
| **Programming language or environment** | R | R | R | R | Python | R | Python | Deep-learning using Keras with the Theano backend on GPUs |

Tables S2, S3, and S4 are spreadsheets saved as comma-separated values (CSV) files and have been uploaded separately.

#### Supplementary figures

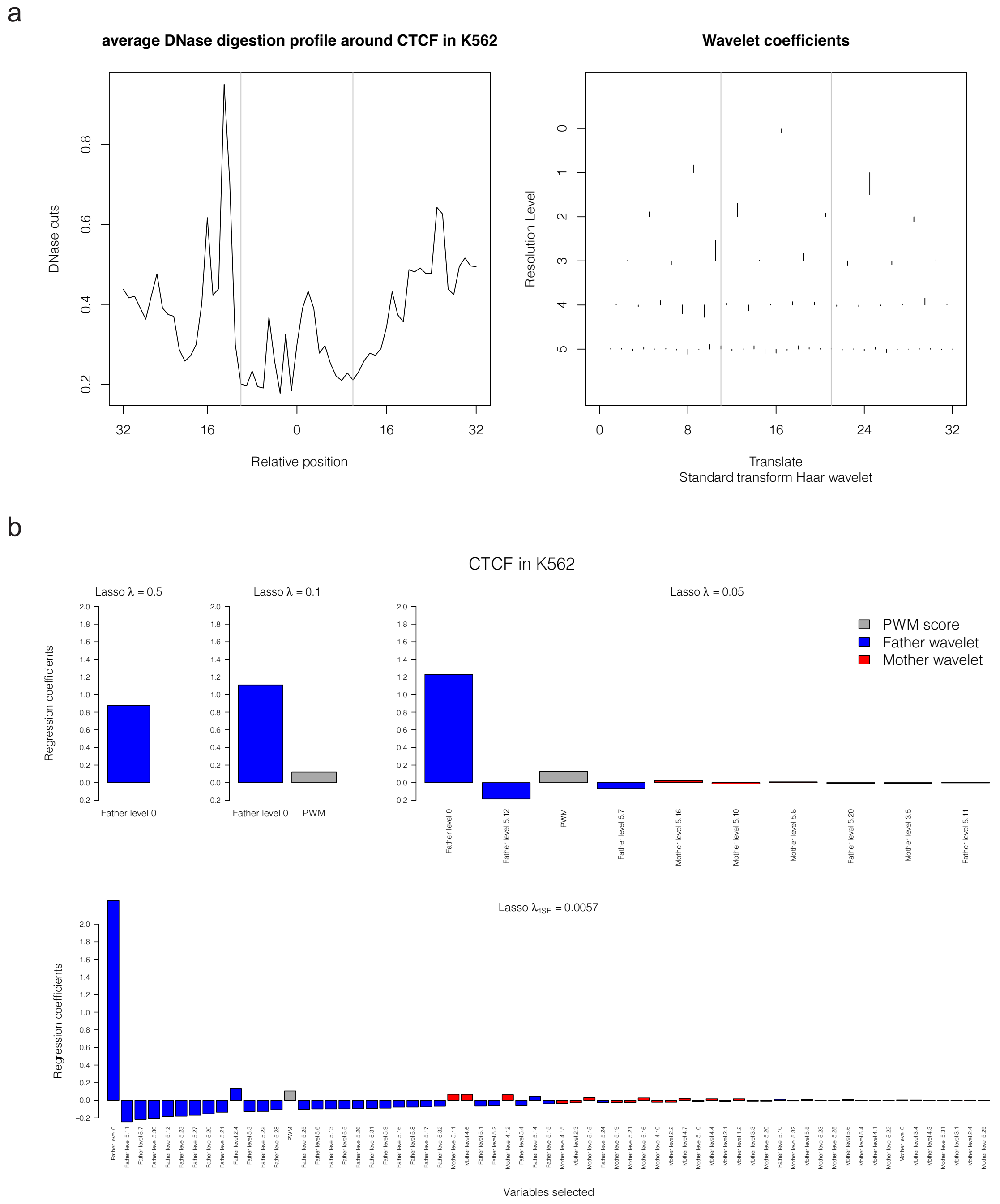


**Fig. S1. Details of wavelet decomposition of CTCF DNase digestion profiles.** (**a**) Example DNase digestion profile and Haar wavelet coefficients. Left: average DNase digestion profiles around CTCF motif matches in K562 cell type in a window of size 64 bp. Right: Haar wavelet coefficients at different resolution levels for those DNase digestion profiles. (**b**) Selecting regression coefficients for CTCF in K562 cell line using Lasso with $\lambda$ equals 0.5, 0.1, 0.05, or $\lambda_{1SE}$ (largest value of $\lambda$ such that mean cross-validated error is within one standard error of the minimum). Selected variables are ordered by the absolute value of coefficients. PWM score is in gray, mother wavelet coefficients are in red, and log2 transformed father wavelet coefficients are in blue. The names of the wavelet coefficients (bars) show the location indices of the wavelet coefficients (e.g. 'Father level 5.12' refers to the 12th father wavelet coefficient at resolution level 5).

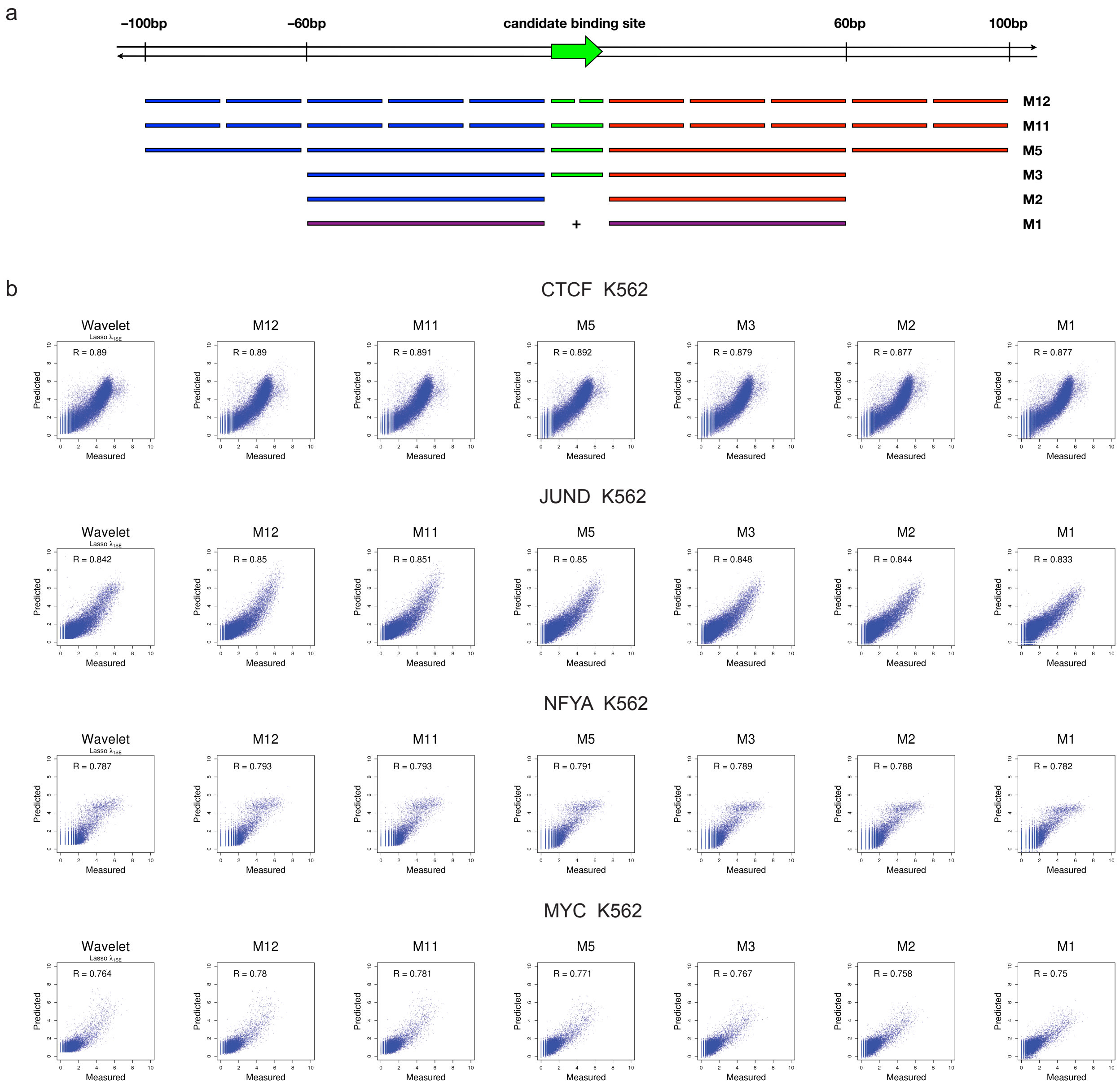


**Fig. S2. Details of MILLIPEDE binning and performance in comparison with wavelet models.** (**a**) Understanding the relationships between bins in various MILLIPEDE models. All bins are defined relative to the strand orientation of the candidate binding site: green bins are within the binding site, blue bins are upstream, red bins are downstream, and purple bins in the M1 model sum together cleavage events from the M2 model's blue and red bins. Models are arranged from most to least complex [1]. In this work, we use model M5, which has two upstream bins, a bin spanning the motif site, and two downstream bins. (**b**) Examples of prediction performances in 5-fold cross-validation using DNase wavelet coefficients and different MILLIPEDE bins. The wavelet models used variables selected using Lasso with $\lambda_{1SE}$ (largest value of $\lambda$ such that mean cross-validated error is within one standard error of the minimum). The M5 model outperforms the wavelet model in all these examples.

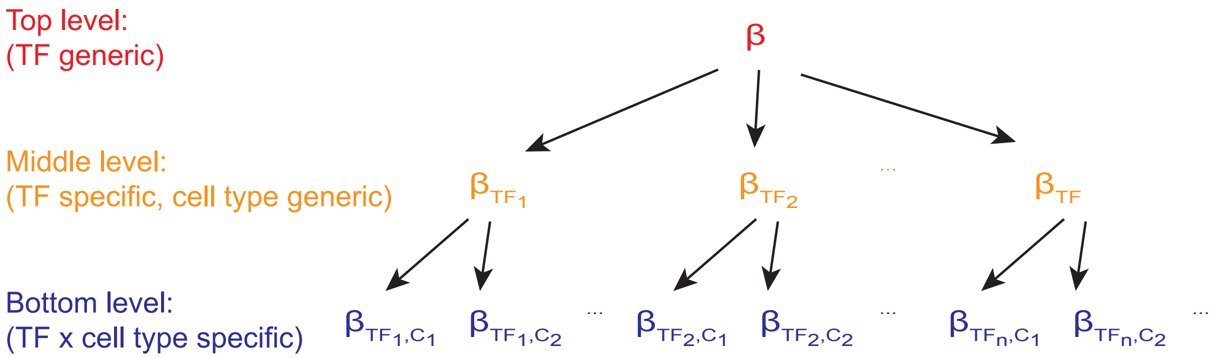


**Fig. S3. Bayesian hierarchical model structure.** Model parameters are organized in a hierarchy to allow for borrowing or sharing of information. At the bottom level are regression parameters specific to a particular TF × cell-type combination. For each TF, the bottom level parameters are themselves drawn from a shared distribution for the TF at the middle level. Likewise, the parameters associated with each TF's distribution at the middle level are themselves drawn from a single shared distribution for all TFs at the top level.

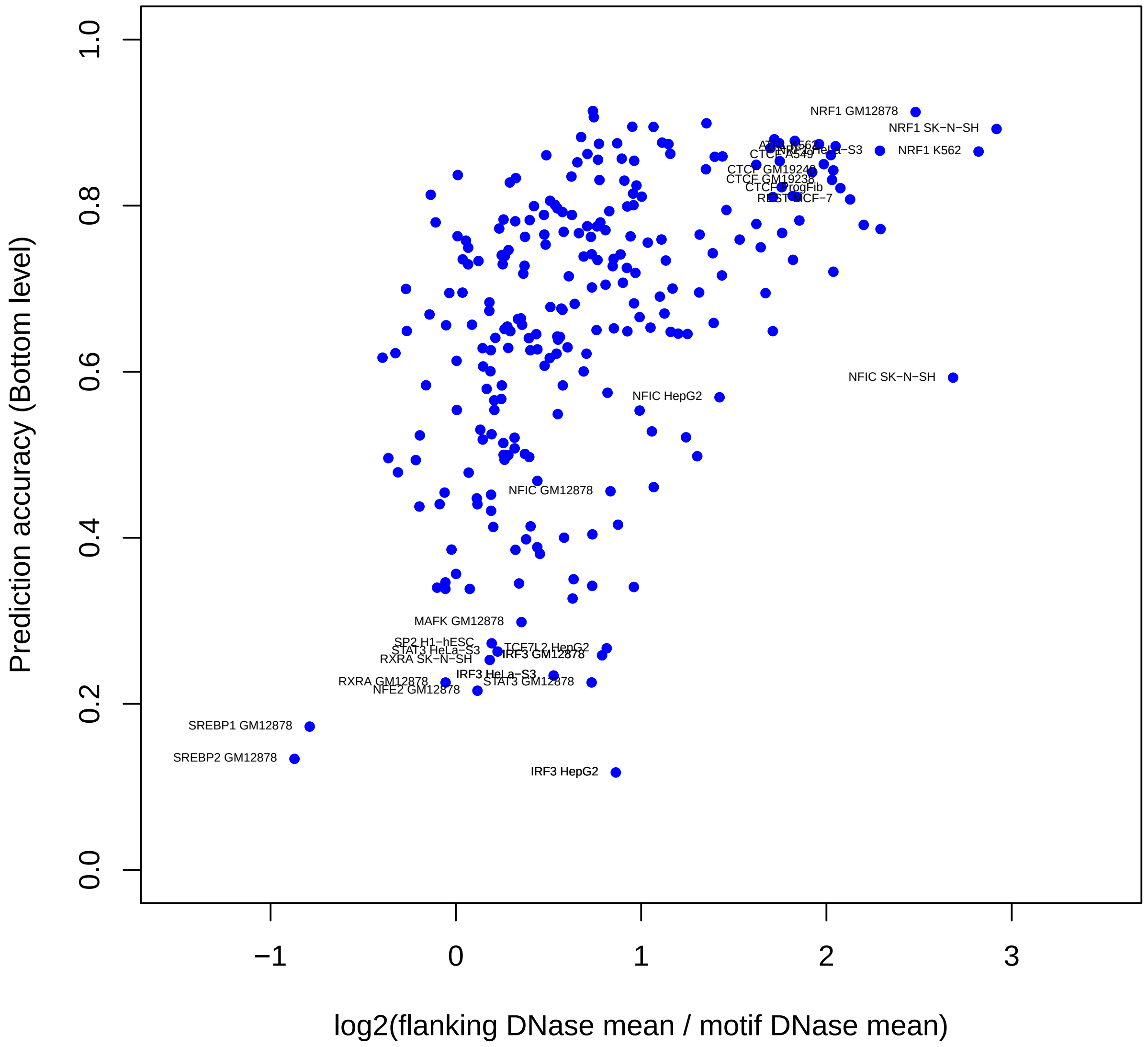


**Fig. S4. Prediction accuracy (bottom level) as a function of DNase depletion ratio.** DNase depletion ratio was calculated as the log ratio of the average number of DNase cleavage events in the 60 bp proximal flanking regions divided by the average number of DNase cleavage events within the motif itself. Each dot represents one TF × cell-type combination in the Duke DNase data.

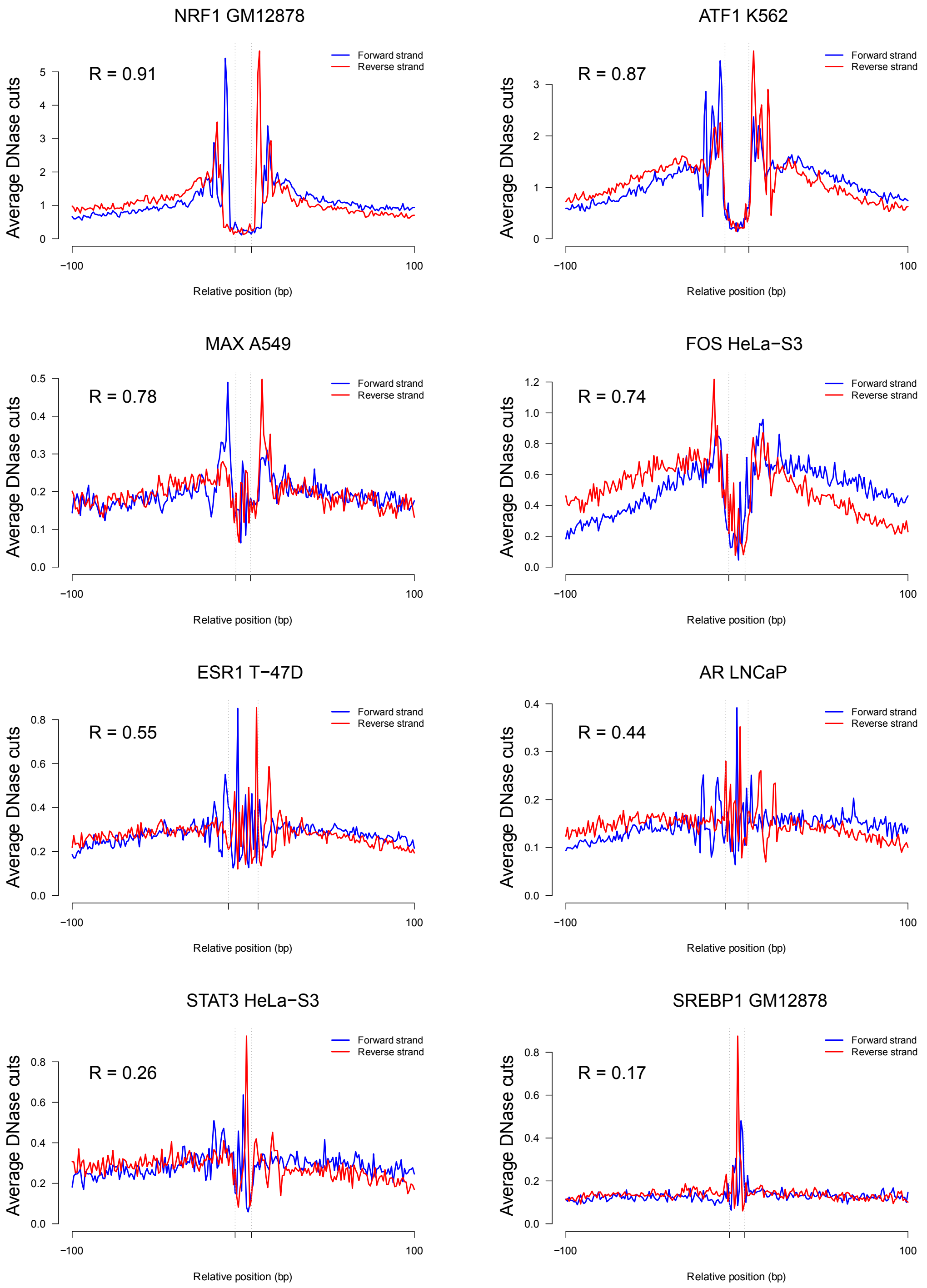


**Fig. S5. Examples of DNase digestion profiles on both strands around a motif, showing TF × cell-type combinations with high to low prediction accuracy.** DNase profiles were averaged over 1000 binding sites with the highest occupancy measured by ChIP-seq. The TF × cell-type combinations that achieved higher prediction accuracy tend to show much clearer DNase depletion patterns (depletion of DNase cleavage events within the motif region, coupled with elevation in the proximal flanking regions).

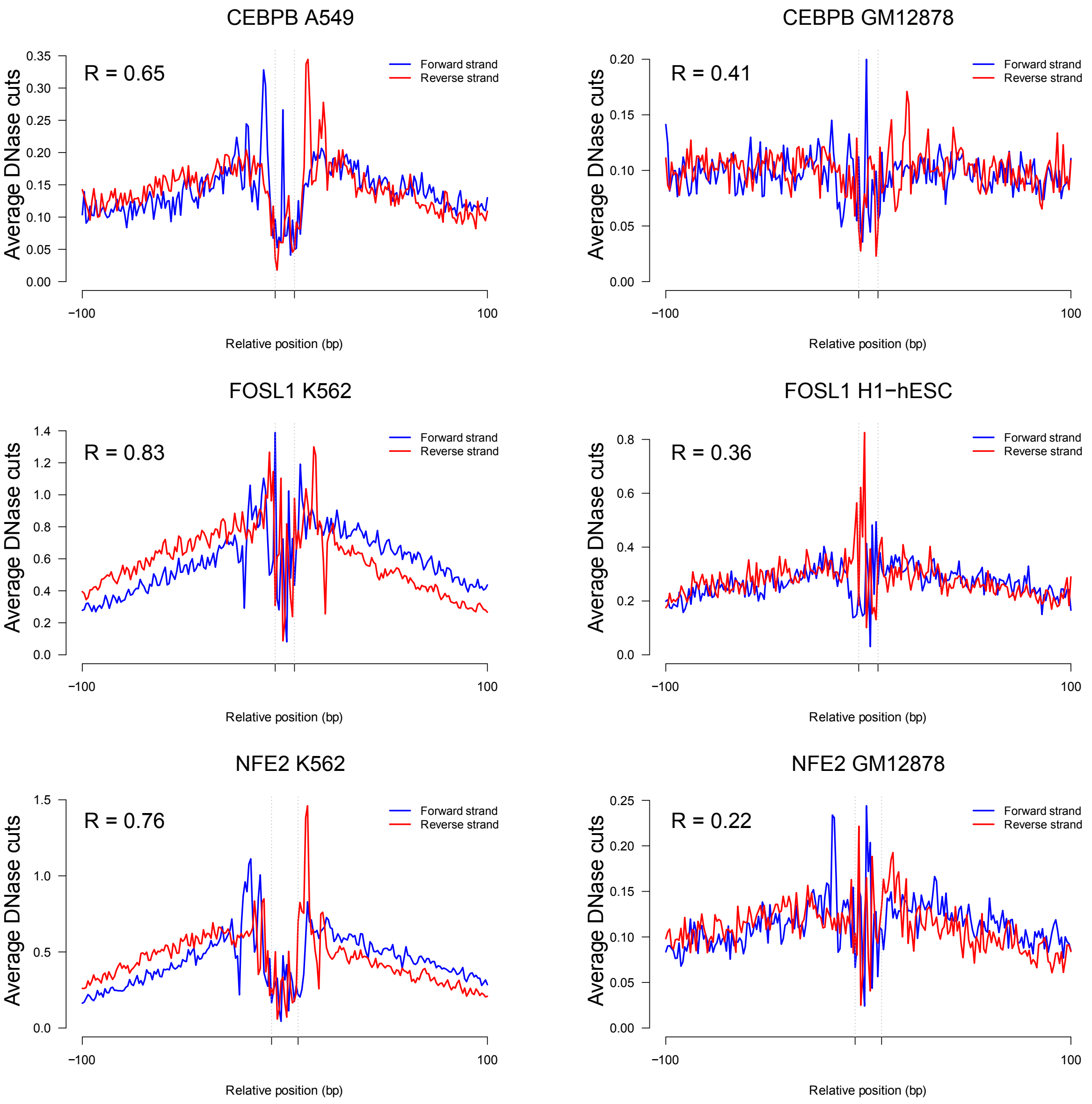


**Fig. S6. Examples of DNase digestion profiles for TF × cell-type combinations with higher prediction accuracy in one cell type (left) but lower prediction accuracy in a different cell type (right).** Note that scales on left and right differ. Consistent with fig. S5, cell types with higher prediction accuracy often show much clearer DNase depletion patterns.

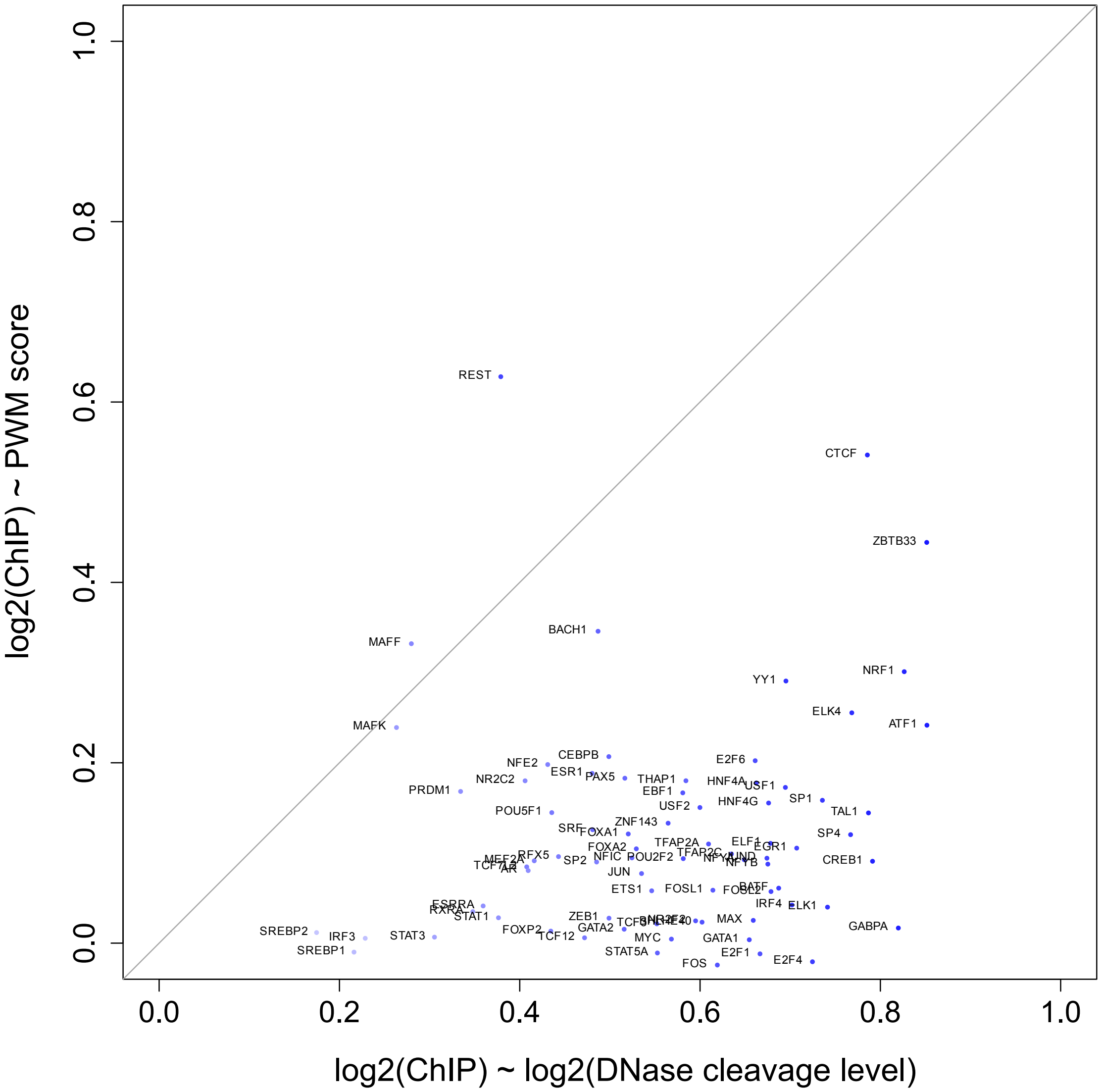


**Fig. S7. Scatter plot comparing the correlation of measured TF occupancy with total number of nearby DNase cleavage events vs. the correlation of measured TF occupancy with PWM score.** X-axis: Pearson correlation of log2(measured TF occupancy) with log2(total number of nearby DNase cleavage events). Y-axis: Pearson correlation of log2(measured TF occupancy) with PWM score. Overall level of nearby DNase cleavage is more correlated with measured TF occupancy than PWM score is—and typically markedly so—for nearly all TFs, with REST and MAFF being exceptions.

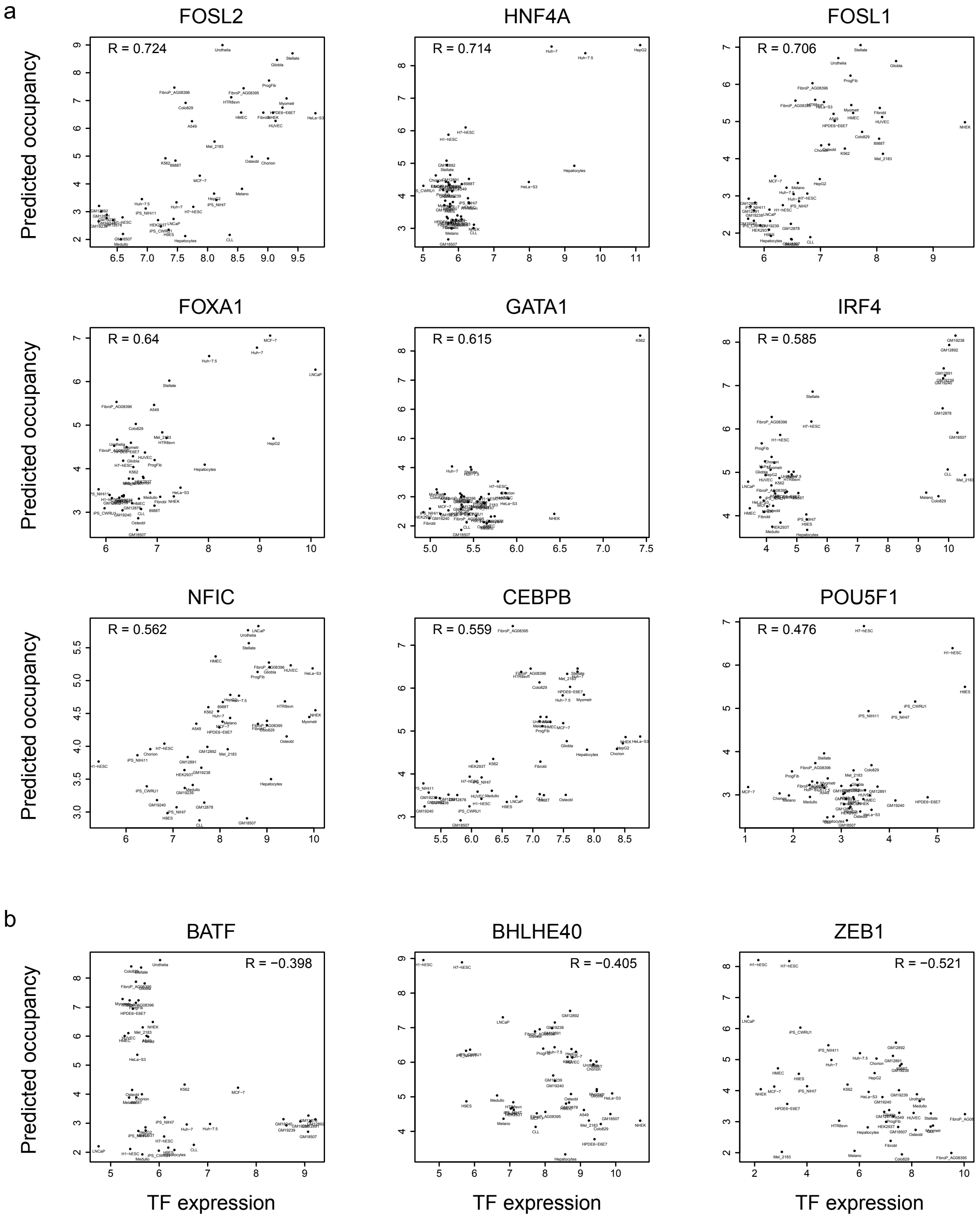


**Fig. S8. Examples showing strong relationships between TF occupancy and TF expression in different cell types.** (**a**) The nine TFs exhibiting the strongest positive correlation between predicted TF occupancy and measured TF expression level (which serves as a rough but imperfect proxy for active nuclear TF concentration). (**b**) The three TFs exhibiting statistically significant negative correlation between predicted TF occupancy and measured TF expression level. Interestingly, BATF has high expression but low predicted occupancy in lymphoblastoid cells, while IRF4, shown in (a), has high expression and high predicted occupancy in lymphoblastoid cells.

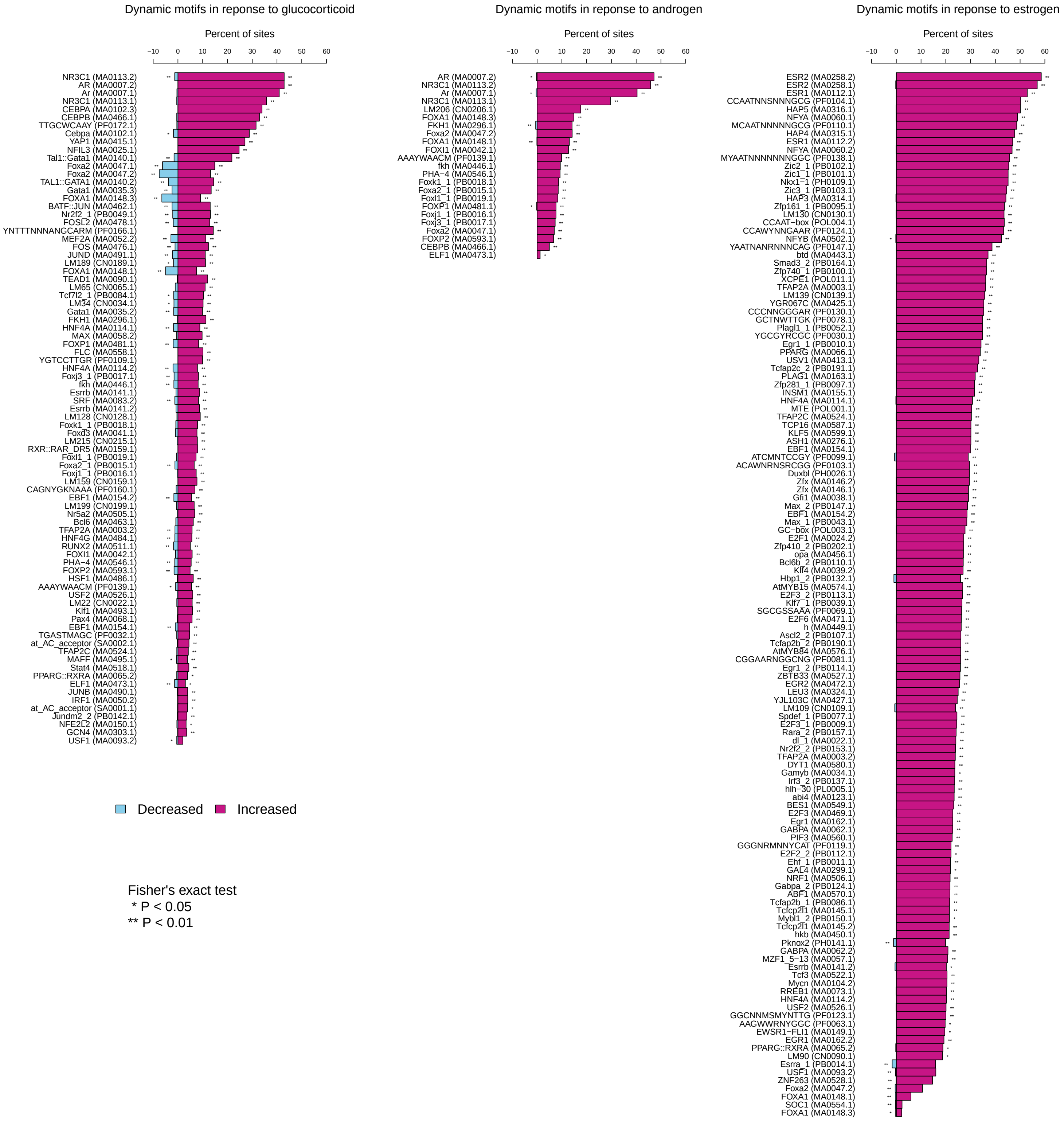


**Fig. S9. TF occupancy dynamics in response to hormone stimulation.** The three panels are analogous to the ones shown in Fig. 5 of the manuscript, showing the same analysis but here conducted at the level of individual TFs rather than RSAT TF motif clusters. Motif identifiers are displayed in parentheses next to motif names.

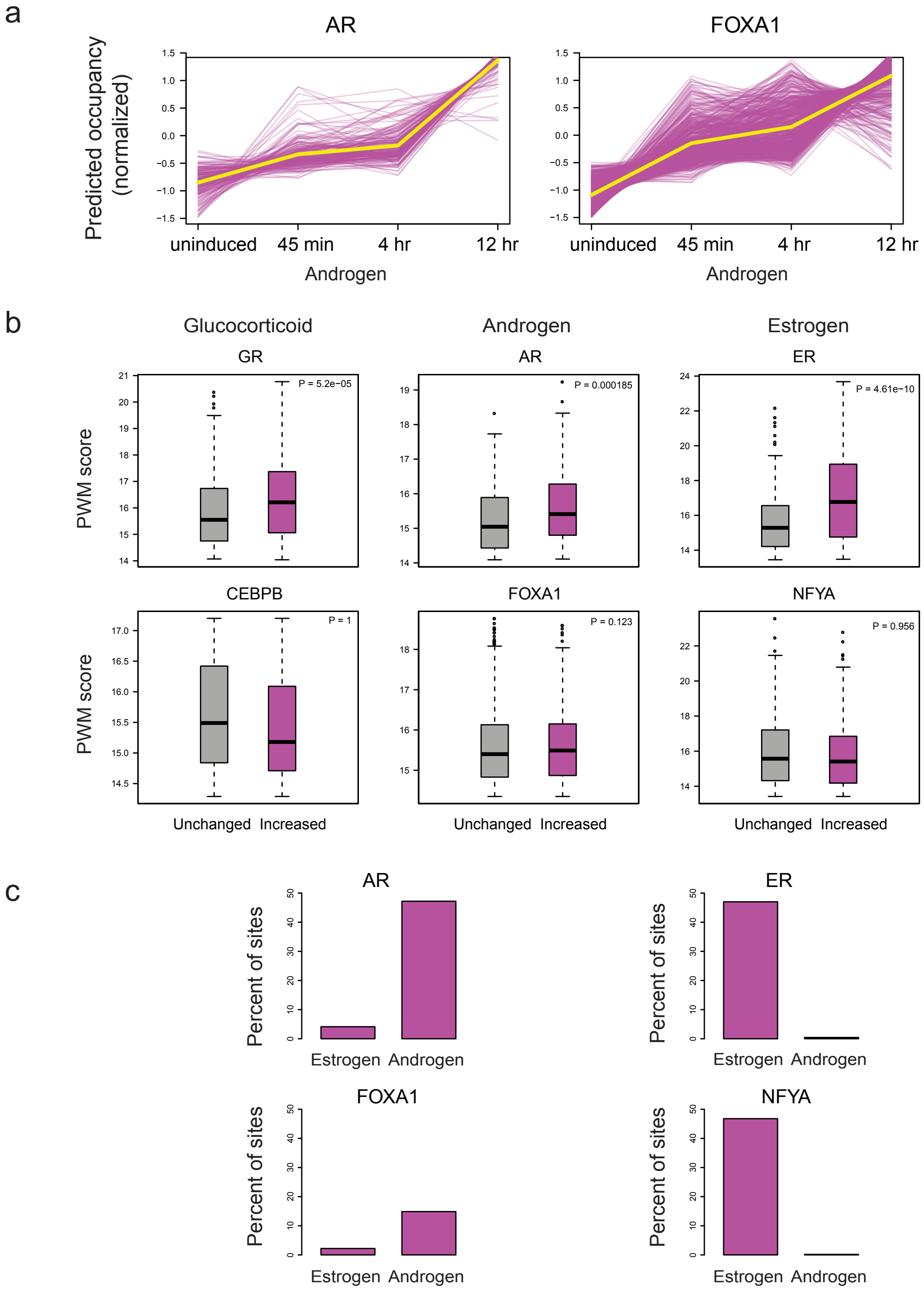


**Fig. S10. Further details of TF occupancy dynamics in response to hormone stimulation.** (**a**) Under androgen stimulation, AR and FOXA1 sites increase in occupancy gradually over the time course, revealing the importance of a quantitative perspective on occupancy. (**b**) For GR in glucocorticoid stimulation, AR in androgen stimulation, and ER in estrogen stimulation, sites with significantly increased occupancy possess significantly higher average PWM scores than sites with unchanged occupancy. This is not true for CEBPB, FOXA1, or NFYA, the second-most responsive TFs in the respective treatment conditions. (**c**) Specificity of increased occupancy. Bar plots on the left show the response of AR and FOXA1, revealing that their increased occupancy is highly specific to androgen induction. Bar plots on the right show the response of ER and NFYA, revealing that their increased occupancy is highly specific to estrogen induction.

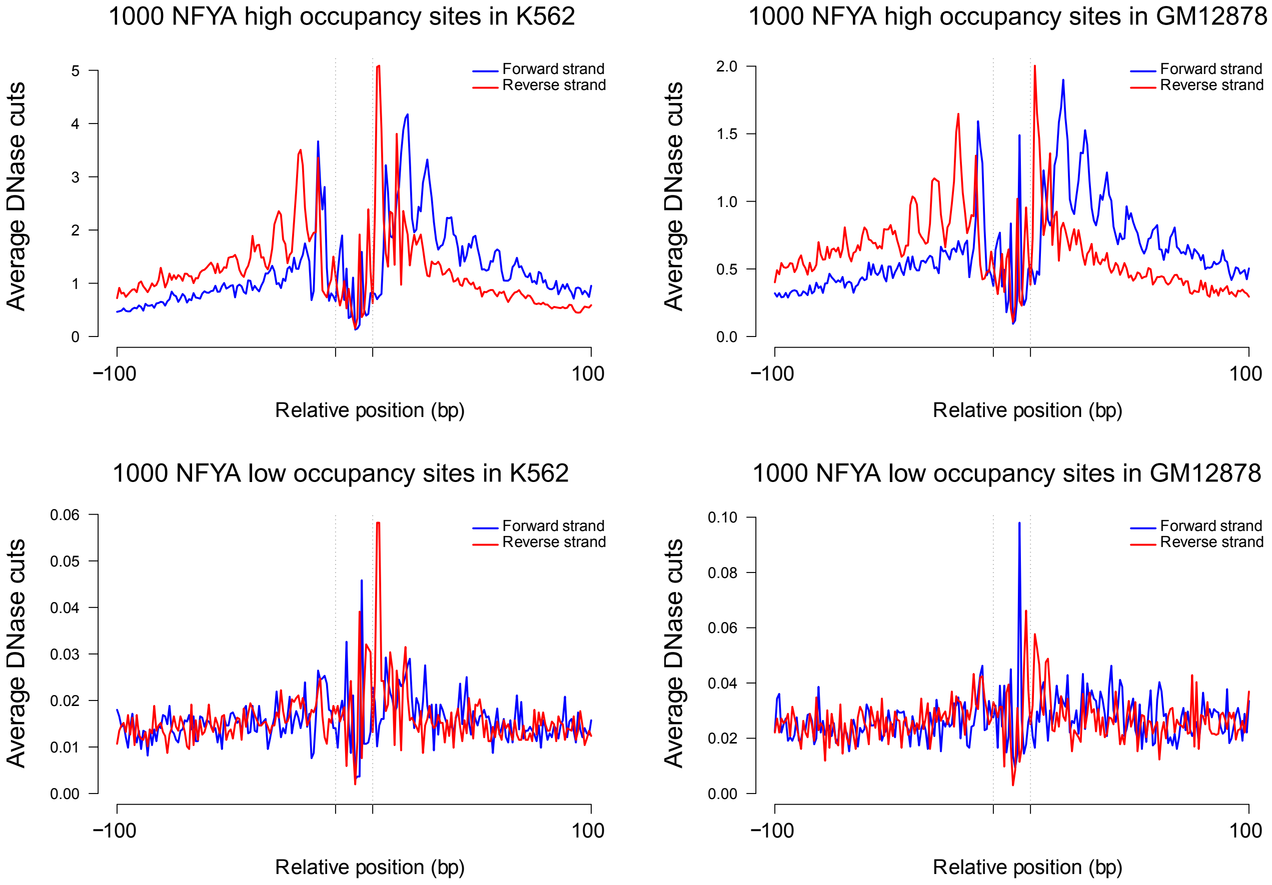


**Fig. S11. Average DNase digestion profiles around 1000 high occupancy and 1000 low occupancy NFYA sites in K562 and GM12878 cell types.** In both cell types, the oscillation patterns of DNase cleavage events in the flanking regions of NFYA high occupancy sites (top row) are similar to the DNase oscillation patterns previously observed within nucleosomes [2], suggesting that NFYA is perhaps more likely to bind flanked by nucleosomes.

a

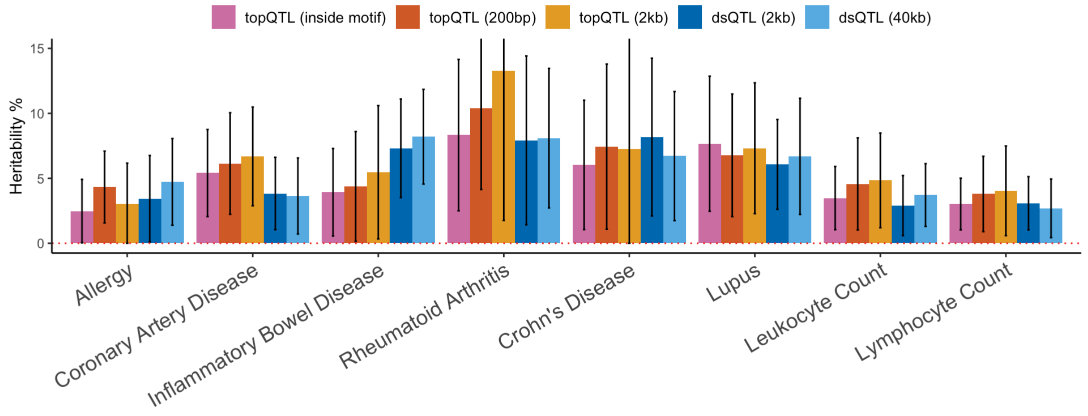


b

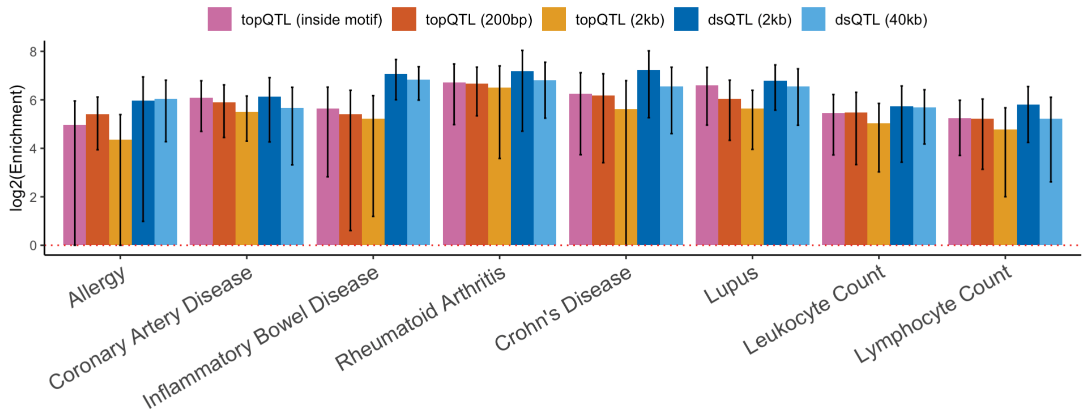


**Fig. S12. Heritability and enrichment estimates for topQTLs and dsQTLs.** Heritability (**a**) and log2(enrichment) estimates (**b**) for topQTLs and dsQTLs in different diseases and complex traits using S-LDSC. Lead SNPs for both topQTLs and dsQTLs were used as binary annotations. Bars represent topQTLs from SNPs within motif matches, topQTLs within 200bp and 2kb around motif matches, and dsQTLs within 2kb and 4kb of the 100 bp DNase window. Error bars represent 95% confidence intervals and were truncated at 15% in the heritability figure.

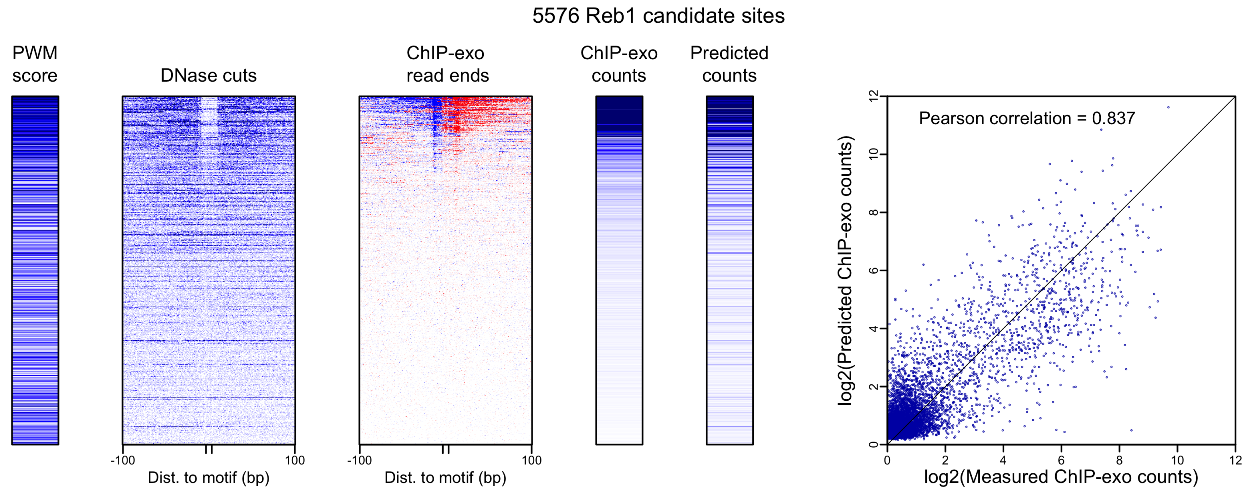


**Fig. S13. Predicting quantitative Reb1 occupancy in yeast.** (Non-hierarchical) regression model was trained on ChIP-exo read counts around Reb1 candidate binding sites in yeast. Rows in the left panels correspond to candidate binding sites and were ordered by the measured number of ChIP-exo read counts (column 4). Darker colors mean higher PWM score, higher number of DNase cleavage events, or higher occupancy (ChIP-exo read counts).

### References

[1] Luo, K. & Hartemink, A. J. Using DNase digestion data to accurately identify transcription factor binding sites. In *Pac. Symp. Biocomputing*, 80–91 (World Scientific, Hackensack, NJ, 2013).

[2] Zhong, J. *et al.* Mapping nucleosome positions using DNase-seq. *Genome Res.* **26**, 351–364 (2016).
